# A chromatinized origin reduces the mobility of ORC and MCM through interactions and spatial constraint

**DOI:** 10.1101/2023.05.18.541303

**Authors:** Humberto Sánchez, Zhaowei Liu, Edo van Veen, Theo van Laar, John F. X. Diffley, Nynke H. Dekker

## Abstract

Chromatin replication involves the assembly and activity of the replisome within the nucleosomal landscape. At the core of the replisome is the Mcm2-7 complex (MCM), which is loaded onto DNA after binding to the Origin Recognition Complex (ORC). In yeast, ORC is a dynamic protein that diffuses rapidly along DNA, unless halted by origin recognition sequences. However, less is known about the dynamics of ORC proteins in the presence of nucleosomes and attendant consequences for MCM loading. To address this, we harnessed an *in vitro* single-molecule approach to interrogate a chromatinized origin of replication. We find that ORC binds the origin of replication with similar efficiency independently of whether the origin is chromatinized, despite ORC mobility being reduced by the presence of nucleosomes. Recruitment of MCM also proceeds efficiently on a chromatinized origin, but subsequent movement of MCM away from the origin is severely constrained. These findings suggest that chromatinized origins in yeast are essential for the local retention of MCM, which may facilitate subsequent assembly of the replisome.

## INTRODUCTION

Eukaryotic genomes are structurally organized as a combination of DNA and proteins known as chromatin. Cell biology, genetics, and biochemical experiments have demonstrated that the quantity and local density of the basic structural unit of chromatin, the nucleosome, can define where the initiation of DNA replication happens and that chromatin folding influences replication timing^1^. Less is known about the influence of nucleosomes on the individual proteins that are involved in establishing DNA replication, including their attendant dynamics.

DNA replication in eukaryotes occurs during the S phase of the cell cycle, during which the entire genome must be duplicated in a coordinated manner. In the budding yeast *Saccharomyces cerevisiae*, replication is initiated from hundreds of sequence-specific origins of replication. Each origin is prepared for replication initiation in two temporally separated steps that are essential for proper DNA replication^2^. During the G1 phase of the cell cycle, the Origin Recognition Complex (ORC) first binds to replication origins^3^. These origins are autonomous replicating sequences (ARS) that contain both a strong binding site (the ARS consensus sequence, or ACS) and a weak binding site (B2) for ORC. Once bound, ORC, together with Cdc6 and Cdt1, then loads two inactive Mcm2-7 (MCM) replicative DNA helicases, forming the pre-replication complex (pre-RC); this is known as the licensing step^4, 5^. During S phase, origins of replication are activated by the combined action of two kinases, S-CDK and DDK, and several other proteins known as firing factors, which serve to bring Cdc45 and GINS into association with MCM and form the CMG holo-helicase^6–8^. The addition of DNA polymerases as well as accessory proteins completes the assembly of the full replisome and allows DNA replication to proceed^8, 9^.

During the licensing step, ORC, Cdc6 and MCM/Cdt1 load at DNA replication origins that are embedded in the chromatin^2^. Biochemical studies have shown that such origins are free of nucleosomes^10^, and genome-wide analysis suggests that these specific nucleosome-free regions (NFRs) favor ORC binding^11^. Interestingly, ORC then plays an active role in positioning nucleosomes around the origin^11, 12^ by coordinating the activity of chromatin remodelers before and during replication^12, 13^. Chromatin remodelers that act at the origin also influence origin licensing^14^.

We have previously demonstrated that yeast ORC is a mobile protein that rapidly diffuses on bare DNA, but that origin recognition halts this search process^15^. Furthermore, it has been observed that single and double MCM hexamers diffuse on DNA^4, 5^ and that nucleosomes can act as potential obstacles for MCM diffusion^16^, so we sought to investigate the roles of origin-flanking nucleosomes in either directly limiting or locally targeting the diffusion of ORC and, consequently, in the loading of MCM onto DNA.

## RESULTS

### Establishment of a labeled chromatinized origin of replication within 10.4 kbp DNA for single-molecule investigations

To experimentally examine the origin localization and dynamics of ORC and MCM on a chromatinized origin of replication, we designed an *in vitro* single-molecule force-fluorescence assay using purified yeast proteins (**Supplementary Figure 1.1**). We wished to visualize the chromatinized origin using yeast histone octamers marked with fluorescent labels. To achieve this, we first introduced a single cysteine on the H2A histone by replacing residue lysine 120 (K120), which avoids disruption of the overall nucleosome structure (**Figure 1a**; Methods). Purified yeast histone octamers containing two H2A (K120C) histones were then covalently bound with one single AF488 dye per cysteine (degree of labeling *p_label_* = 0.81, as determined from bulk experiments; Methods). We next engineered an ARS1 origin of replication flanked by two nucleosome positioning sequence (NPS) sites spaced by 150 bp (50 nm) (Ref. 16 and Methods), and reconstituted fluorescently labeled nucleosomes onto these NPS sites using salt gradient dialysis. The resulting chromatinized origin tested positively for *Pst*I restriction digestion between the nucleosomes (**Supplementary Figure 1.2b**), which indicated that it should remain accessible to ORC and MCM. Next, we ligated this chromatinized origin to two biotinylated DNA fragments of distinct sizes. This resulted in a 10.4 kbp chromatinized DNA molecule (**Figure 1b**) in which the origin was localized at approximately one third of the total length of the DNA molecule (**Supplementary Figure 1.2c**). In addition to the ARS1 origin, the 10.4 kbp DNA molecule contained a number of endogenous potential binding sites for ORC^17^ and regions of high AT content (**Supplementary Figure 1.2d**).

**Figure 1.**
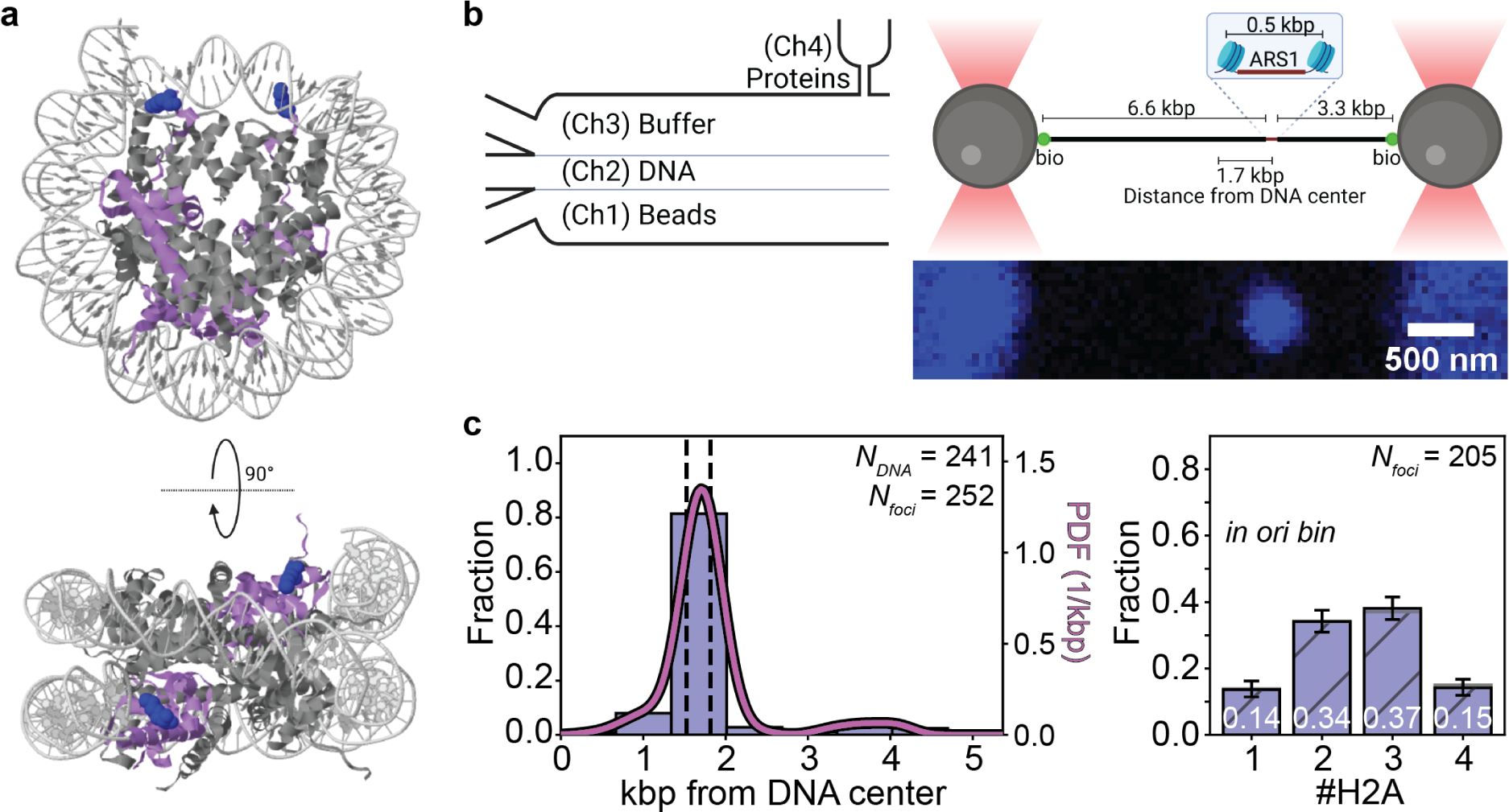
Design and characterization of a labeled chromatinized origin within 10.4 kbp DNA. **(a)** Structural depictions of the yeast nucleosome (PDB: 1ID3), including in blue the mutated residues on the H2A histones for adding fluorescent labels. **(b)** (left panel) Schematic of the flow cell used in the single-molecule experiments. (right upper panel) Schematic of the 10.4 kb DNA held in an optical trap that contains the ARS1 origin of replication flanked by two nucleosome positioning sites (NPSs). The DNA is chromatinized via salt gradient dialysis prior to its introduction into the single-molecule flow cell (**Supplementary Figure 1.2**). (right lower panel) Confocal scan showing that signal from fluorescently labeled H2A histones is detected as a single diffraction-limited spot localized at the NPSs. **(c)** (left panel) Spatial distribution of fluorescently labeled H2A on DNA described in panel (b), as deduced from the blue diffraction-limited spots (*N_foci_*) collected from 248 distinct DNA molecules (*N_DNA_*). Dashed lines indicate the location of the NPSs, and the solid curve indicates the kernel density estimation of the data. (right panel) Stoichiometry distribution of fluorescently labeled H2A histones in the bin containing the chromatinized origin. Error bars are one-sigma Wilson confidence intervals. White numbers inside the bars designate the fitted values based on the model described in the Methods and in **Supplementary Figure 1.3**. Data in both panels derives from four chromatinized samples (**Supplementary Figure 1.4**).

Following preparation, the 10.4 kbp DNA molecules including a chromatinized origin were introduced into the microfluidic flow cell of our single-molecule instrument. Here, the DNA molecules were tethered to streptavidin-coated beads in a dual-beam optical trap, after which they were visualized in a protein-free channel of the flow cell (buffer channel) while under a tension of 2 pN. Nucleosome-bound AF488-H2A was detected as a bright fluorescent spot (or focus) whose spatial position was determined (**Figure 1b**). By repeating this measurement on multiple DNA molecules, we could build up the spatial distribution of H2A foci as a function of genomic coordinate. As each individual DNA molecule was tethered in one of two possible orientations in the dual-beam optical trap, we report spatial distributions from the midpoint of the DNA. Each histogram bin (width 0.670 kbp; Methods) thus contains the average of the occupancies of two segments of DNA located symmetrically about the midpoint of the DNA: for example, the origin bin contains the mean of the occupancies of the actual origin- and NPS-containing segment and the DNA segment located oppositely from the midpoint of the DNA by 1.7 kbp. The spatial distribution of H2A fluorescent foci (filtered to remove foci containing more than 4 H2A, which could represent aggregates; Methods) showed a clear peak in the bin containing the NPS sites (left panel in **Figure 1c**), suggesting preferential nucleosome assembly at the NPS sites. This spatial distribution of H2A was built up from distinct DNA samples chromatinized on different days, which exhibited only minimal differences between them (**Supplementary Figure 1.4**), highlighting the reproducibility of our sample preparation.

Ideally, our chromatinized origin should contain two fluorescently labeled nucleosomes within a single diffraction-limited spot. Assuming that the identification of a single fluorescently labeled H2A reflects the presence of a single H2A-H2B dimer bound to a H3-H4 tetramer, we should then measure 4 fluorescently labeled H2A per focus. However, the experimental counts of fluorescently labeled H2A may be reduced by non-unity values for the probability of occupancy of the two NPSs (*p_occupancy_*), the probability of H2A-H2B dimer association to the H3-H4 tetramer (*p_h2a_*), and *p_label_* (**Supplementary Figure 1.3a**). To count the number of labeled H2A, we identified step-wise photobleaching events using Change-Point Analysis (CPA; Methods), whereby the methodology is assessed and validated using dCas9 tagged and labeled identically to H2A (**Supplementary Figure 1.1**). The lifetime prior to photobleaching of AF488 when linked to H2A was similar to that measured when it was linked to dCas9, indicating that labeled H2A are stable on the DNA during the experiment (**Supplementary Figure 1.3c**). Using this photobleaching approach, we found that the H2A foci primarily contained two to three AF488-labeled H2A histones, as shown in the stoichiometry distribution (right panel in **Figure 1c**) built up from a total of *N_DNA_* = 247 derived from the four different chromatinized DNA samples mentioned above (again exhibiting only minimal differences between them, **Supplementary Figure 1.4**). With the independently measured *p_label_* = 0.81 as fixed parameter, the best fit to the experimentally determined stoichiometry distribution (**Figure 1c**, with mean squared error (MSE) = 5.18 x 10^-5^ and fit values described by white numbers inside the bar plot; **Supplementary Figure 1.3b**) yielded *p_occupancy_* = 1.00 (corresponding to full occupancy of the NPSs) and *p_h2a_* = 0.77.

To confirm that the detected fluorescent H2A foci reflected embedding of H2A into nucleosomes – as opposed to unspecific electrostatic interactions of H2A-H2B dimers with the DNA^18^ – we separately probed for nucleosome presence on our 10.4 kbp DNA using force spectroscopy. Previous experiments have shown that under force, DNA can be unwrapped from the histone octamer. Irreversible jumps in the force-extension curve in which 27 nm (80 bp) of DNA is unwrapped from either a hexasome or a full nucleosome have been reported to occur over a broad range of forces (8-40 pN) ^19–22^. Indeed, when we pulled on one extremity of a tethered chromatinized DNA molecule with labeled H2A histones at a constant speed (100 nm/s) (**Supplementary Figure 1.5a**), we could identify such jumps (**Supplementary Figure 1.5b**). These occurred at a force of 17.6 pN ± 5.8 pN (mean ± standard deviation) and yielded contour length increments of 24.8 nm ± 5.9 nm (mean ± standard deviation), the latter corresponding to the contour length of the unwrapped DNA. At least two such jumps were revealed in 88.5% of the chromatinized DNA molecules probed (**Supplementary Figure 1.5c**), implying that our chromatinized DNA typically contains two nucleosomes (or hexasomes).

Together, these single-molecule fluorescence and force spectroscopy results support our establishment of a 10.4 kbp DNA containing a single origin of replication flanked by two nucleosomes. As the force spectroscopy does not report on nucleosome location, however, to investigate the influence of a chromatinized origin of replication on ORC and MCM, we focused on fluorescence readouts alone.

### Nucleosomes enhance intrinsic ORC preference for the origin of replication

To study the effect of chromatinized origins on ORC binding and dynamics, we labeled the N-terminus of the Orc3 subunit with a JF646 fluorophore via a HaloTag (Methods). We confirmed that the labeled ORC could load MCM in bulk assays (**Supplementary Figure 1b**). We then prepared four different 10.4 kbp DNA molecules that either contained ARS1 or a mutated origin without specific affinity for ORC^15, 23^, and that were chromatinized or not.

We first performed experiments that report on the rapid binding of ORC to a chromatinized origin. To do so, we incubated the optically trapped DNA molecule held at near-zero force in a reservoir of the microfluidic flowcell containing 5 nM JF646-ORC for 5 s. Subsequently, we shifted the DNA to a separate, protein-free channel and imaged it under a stretching force of 2 pN (**Figure 2a**). DNA-bound JF646-ORC could then be observed as a bright fluorescent spot (focus), after which the experiment was repeated with a new DNA. Stoichiometric analysis of the foci via step-wise photobleaching indicated that they predominantly contained individual ORC molecules (**Supplementary Figure 2**). Foci containing more than 5 ORCs, which could represent aggregates (Methods), were not analyzed further. The remaining fluorescent foci were observed throughout the DNA molecule, but the overall spatial distribution exhibited clear overrepresentation of ORC in the bin containing the ARS1 origin (30% of the total, **Figure 2b-i**). This preference for ORC binding in the bin containing the origin^15^ increased to 39% when the origin was chromatinized (**Figure 2b-ii**). Mutation of the origin reduced preferential binding by ORC in the bin containing the origin^15^; instead, ORC was observed to peak in an adjacent bin (**Figure 2b-iii**) that included a potential ORC binding site (located at 2.4 kbp from the start of the construct, **Supplementary Figure 1.2d**). Chromatinization of this mutated origin nonetheless increased ORC presence in the origin bin from 14% to 26% (**Figure 2b-iii, 2b-iv**). For the four conditions probed, ORC binding in the bin containing the origin (chromatinized or not) was subjected to tests of statistical significance (**Figure 2c**). These results indicate a statistically significant enhancement of ORC binding in the presence of either ARS1 or nucleosomes, with the effect of the former being more pronounced.

**Figure 2.**
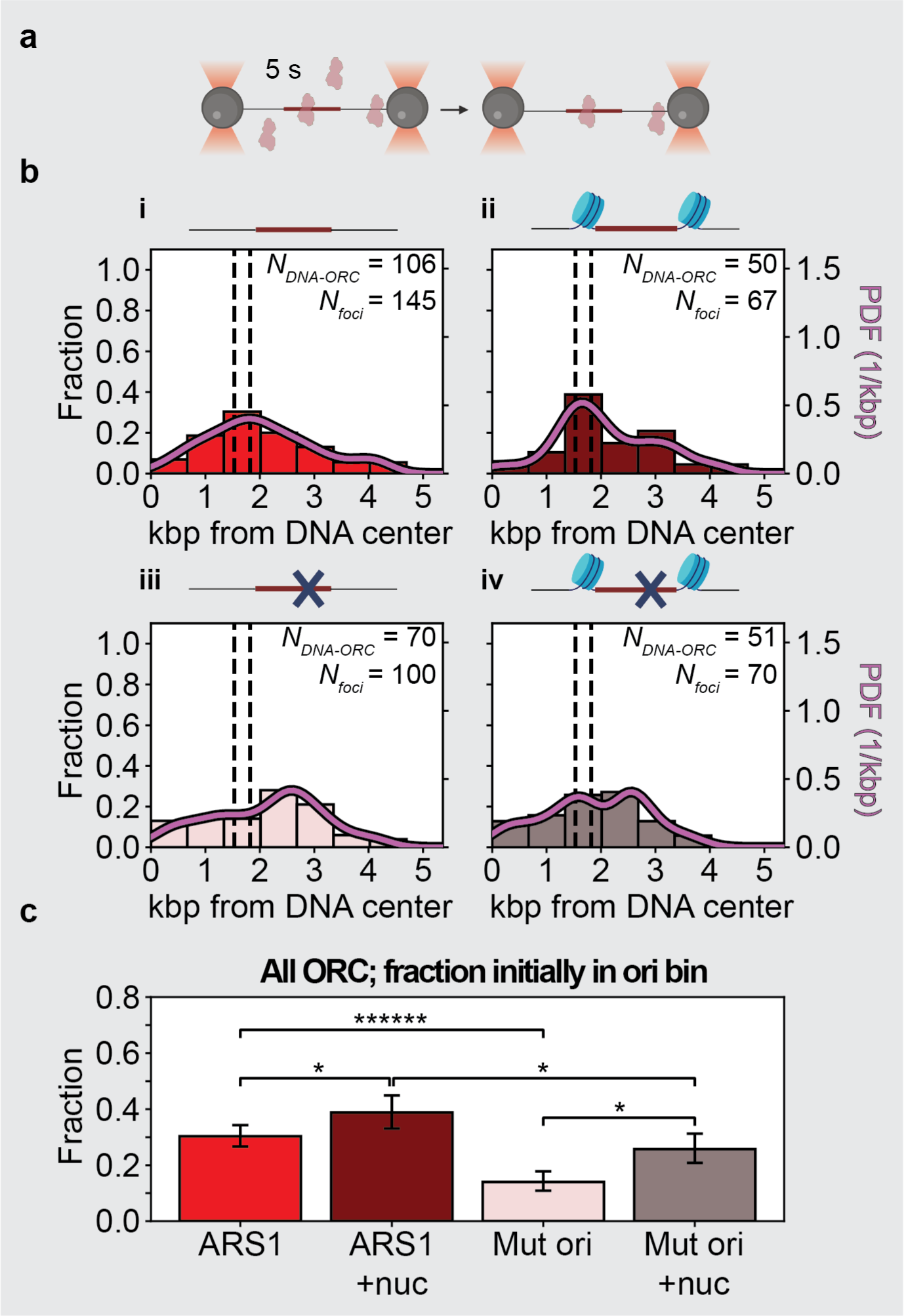
Spatial distribution of rapidly bound ORC on 10.4 kbp DNA containing a chromatinized origin. **(a)** Incubation method pertaining to data in panels (b-c). Tethered DNA molecules (chromatinized or not, as indicated) are introduced into protein channel (Ch4) for incubation with ORC and Cdc6 for 5 s in the presence of ATP, and then moved into the buffer channel (Ch3) for confocal scanning (see Figure 1b for schematic of the flow cell). **(b)** (i) Spatial distribution of ORC on DNA containing a non-chromatinized origin as deduced from the red diffraction-limited spots (*N_foci_*) collected from 106 distinct DNA molecules (*N_DNA-ORC_*). The dashed lines indicate the location of the NPSs, and the solid curve indicates the kernel density estimation of the data. (ii) Spatial distribution of ORC on DNA containing a chromatinized origin as deduced from the red diffraction-limited spots (*N_foci_*) collected from 50 distinct DNA molecules (*N_DNA-ORC_*) analyzed and displayed as in panel (b-i). (iii) Spatial distribution of ORC on DNA containing a non-chromatinized mutated origin as deduced from the diffraction-limited red spots (*N_foci_*) collected from 70 distinct DNA molecules (*N_DNA-ORC_*) analyzed and displayed as in panel (b-i). (iv) Spatial distribution of ORC on DNA containing a chromatinized mutated origin as deduced from the red diffraction-limited spots (*N_foci_*) collected from 51 distinct DNA molecules (*N_DNA-ORC_*) analyzed and displayed as in panel b-i. **(c)** ORC occupancy probability for the bins containing the (chromatinized or not) origin in panels (b-i)-(b-iv). Error bars are one-sigma Wilson confidence intervals. Statistical significance is determined by a two-sided binomial test: * *p* < 0.05, ****** *p* < 0.0000005.

### Nucleosomes flanking the origin reduce ORC mobility and increase ORC lifetime

We have previously shown that ORC is a mobile protein that slides on bare DNA^15^. Here we sought to investigate the motion dynamics of ORC on DNA containing a chromatinized origin following rapid binding under the incubation conditions described above (**Figure 2a**). When we tracked the position of ORC molecules initially located in the bin containing the chromatinized origin over time (**Figure 3a-i**, showing 30% of all traces, selected at random), it appeared that within experimental error many of these ORC molecules hardly changed their position. Conversely, ORC molecules initially located outside the origin (**Figure 3a-ii**, showing 30% of all traces, selected at random) were observed to explore their local DNA environment in seemingly random fashion, which in some cases allowed them to approach, and then move away from, the nucleosomes.

**Figure 3.**
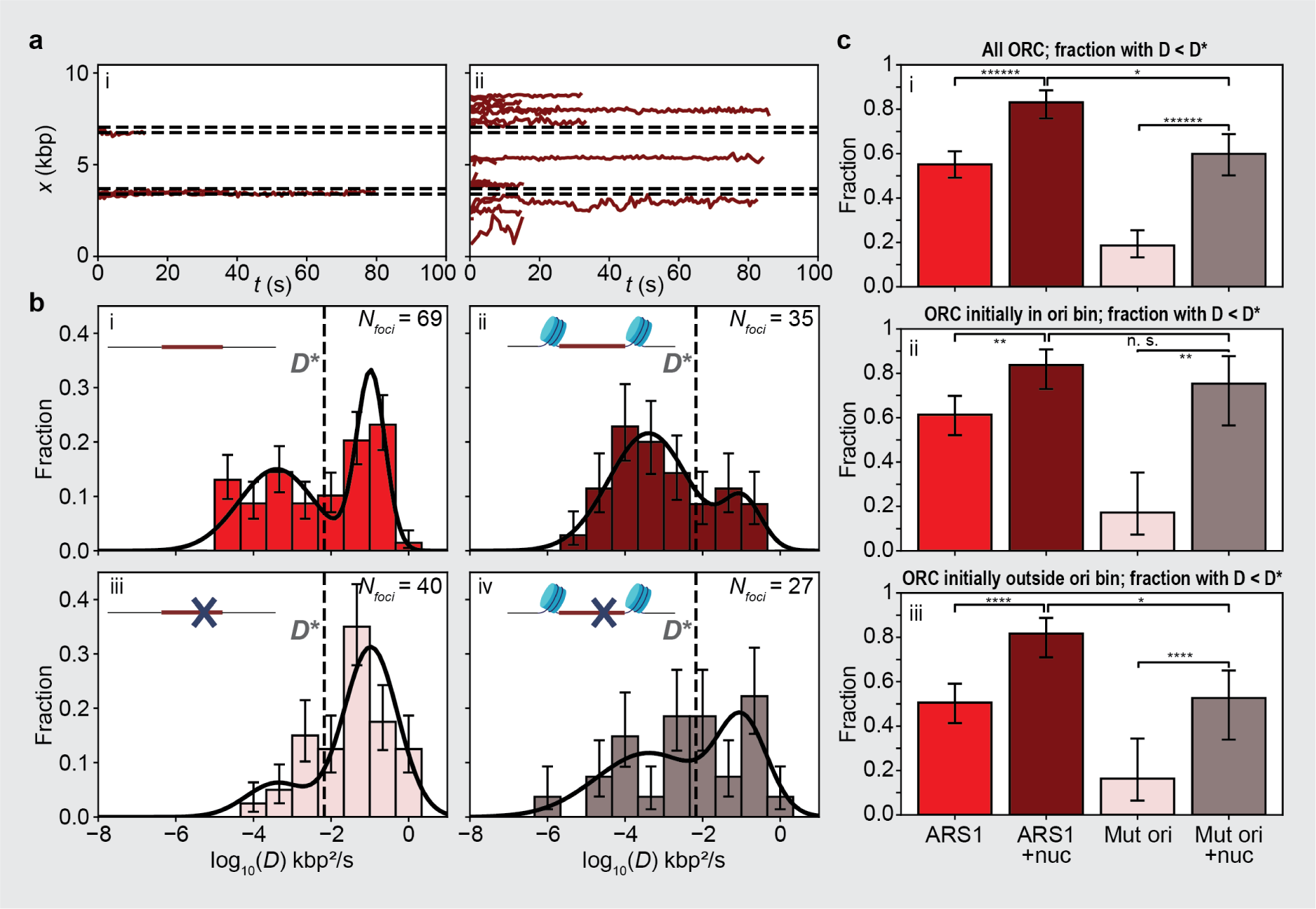
Mobility of ORC on 10.4 kbp DNA containing a chromatinized origin. **(a)** Traces collected on DNA containing a chromatinized origin illustrating motion of JF646-ORC (i) initially localized within the bin containing the chromatinized origin or (ii) initially localized elsewhere. **(b)** Histograms of the diffusion constants of ORC on DNA molecules containing (i) a non-chromatinized origin, (ii) a chromatinized origin, (iii) a non-chromatinized mutated origin, and (iv) a chromatinized mutated origin. Only foci containing 1 or 2 ORC are included in the analysis (*N_foci_*). Error bars are one-sigma Wilson confidence intervals. The observed bimodal distribution is fitted to a double log-normal function (solid black line) which identifies a slowly moving or static ORC subpopulation (55% of the distribution with 0.0049 ± 0.0028 kbp^2^ /s, mean ± SEM) and a fast diffusive ORC subpopulation (0.152 ± 0.023 kbp^2^ /s, mean ± SEM). The dashed line indicates the average of the means of the two normal distributions used to fit log_10_(*D*); this average is used as a threshold (*D*\*). The means of these subpopulations are imposed in the fits to the data in panels (ii-iv). **(c)** Examination of the slowly moving or static ORC subpopulation. (i) Fitted proportion of the slowly moving or static ORC subpopulation for the datasets in panels (b-i)-(b-iv). (ii) The subpopulation of ORC initially localized within the bin containing the (chromatinized or not) origin was extracted from the dataset; the fraction of this subpopulation that is slowly moving or static (*D* < *D\**) is shown here. (iii) The subpopulation of ORC initially localized outside the bin containing the (chromatinized or not) origin was extracted from the dataset; the fraction of this subpopulation that is slowly moving or static (*D* < *D\**) is shown here. Error bars are one-sigma Wilson confidence intervals. Statistical significance is determined by a two-sided binomial test: n. s. not significant, * *p* < 0.05, ** *p* < 0.005, **** *p* < 0.00005, ***** *p* < 0.000005 ****** *p* < 0.0000005.

To quantify these motion dynamics of ORC, we calculated the mean squared displacement for each ORC molecule on the four types of DNA molecules described above versus time interval and extracted a diffusion constant from a linear fit. Distributions (in log scale) of the fitted diffusion constants are shown in **Figure 3b**. We observed a wide spread of diffusion constants, ranging from 10^-5^ to 10^0^ kbp^2^ s^-5^. Using the Bayesian Information Criterion, we identified kinetically distinct population states for ORC on non-chromatinized DNA containing ARS1 (**Figure 3b-i**). These describe either a population with a low mean diffusion constant (hereafter ‘slow ORC population’) (*D_slow_* = 0.0049 ± 0.0028 kbp^2^ s^-1^ (mean ± SEM)) or a population with a higher mean diffusion constant (hereafter ‘fast ORC population’) (*D_fast_* = 0.152 ± 0.023 kbp^2^ s^-5^), consistent with our earlier findings for ORC on longer DNA molecules that contained a synthetic origin of replication^15^. We used these means as imposed values in fitting the distribution of ORC diffusion constants obtained on the other three types of DNA molecules (**Figure 3b-ii, 3b-iii**, and **3b-iv**). As one can observe both by eye and from the fits, chromatinization of the ARS1 origin resulted in an increase in the slow ORC population (**Figure 3b-ii**). On DNA molecules with the mutated origin, the fast ORC population predominated (**Figure 3b-iii**), again consistent with our previous findings^15^ but chromatinization of this mutated origin also resulted in an increase in the slow ORC population (**Figure 3b-iv**). We summarize these results more quantitatively by computing the fraction of the slow ORC population in the total, whereby we used the geometric mean *D\** (= 10^(*μ*slow + *μ*fast)/2^ = 0.0065 kbp^2^ s^-1^, where *μ_i_* are the lognormal population means) as a cutoff (**Figure 3c-i**). This fraction was always observed to be higher upon chromatinization of the origin.

To assess how chromatinization of the origin resulted in an increase of the slow ORC population, we repeated this analysis separately for ORC molecules initially localized in the origin bin (**Figure 3c-ii**) and for ORC molecules initially localized elsewhere (**Figure 3c-iii**). For ORC molecules initially localized in the origin bin, chromatinization of the origin led to an increase in the slow ORC population, irrespective of whether the origin contained ARS1 or was mutated (**Figure 3c-ii**). This reduction in ORC mobility could have resulted solely from the spatial confinement imposed on ORC by the nucleosomes, or additionally through ORC interactions with the nucleosomes. Interestingly, for ORC molecules initially localized elsewhere, chromatinization of the origin also led to an increase in the slow population of ORC molecules, again irrespective of the origin contained ARS1 or was mutated (**Figure 3c-iii**). This reduction in ORC mobility suggested ORC interactions with the nucleosomes.

To address whether chromatinization of the origin influenced the stability of ORC binding, we examined the lifetimes of individual JF646 dyes on DNA-bound ORC molecules by tracking foci containing 1 or 2 ORC molecules at 0.6 s/frame until the fluorescence signal disappeared. Apart from on the DNA molecule with the chromatinized mutated origin, the mean lifetime of ORC was shorter than 30 s, making it substantially shorter than the bleaching-limited mean lifetime of JF646 measured under identical imaging conditions (71.0 s, **Supplementary Figure 1.1c**) by attaching it to DNA-bound dCas9. This indicated that ORC molecules typically dissociated from DNA during the measurement, in accord with our previous observations^15^. Nonetheless, the mean lifetime of the slow ORC population typically exceeded that of the fast ORC population (**Supplementary Figure 3b**). Of the ORC molecules initially located at or close to the ARS1 origin on bare DNA, 61% were associated with the slow ORC population (red bin, **Figure 3c-ii**); this fraction is increased to 84% upon chromatinization of the ARS1 origin (‘bordeaux’ bin, **Figure 3c-ii**). While not all of these ORC molecules may be specifically bound to DNA (as also suggested by a corresponding increase in the slow ORC population observed upon chromatinization of the mutated origin, compare grey and pink bins in **Figure 3c-ii**), nonetheless the increase in the slow ORC population with its higher mean lifetime on a chromatinized ARS1 suggests that flanking nucleosomes can contribute to the temporal retention of ORC near an origin of replication. This could in turn contribute to preferential recruitment of MCM there.

### Stable ORC binding to DNA requires the origin of replication

We also performed experiments with more extended incubation conditions compatible with the timescale required for full maturation of MCM^5, 15^. When performed with ORC and Cdc6 only (see next section for experiments involving MCM), these experiments provided a read-out of long-lived, stable ORC binding. Concretely, we incubated ORC and Cdc6 in bulk, for 30 min with the four types of DNA molecules (untethered to beads, hence with free ends), prior to imaging in single-molecule conditions. We separately verified that an extended incubation period alone did not affect the distributions of H2A position or stoichiometry on chromatinized DNA (**Supplementary Figure 4.1**). We then introduced these pre-incubated DNA molecules into the flowcell, trapped, them, and imaged stable ORC molecules that remained bound to the DNA (**Figure 4a**).

**Figure 4.**
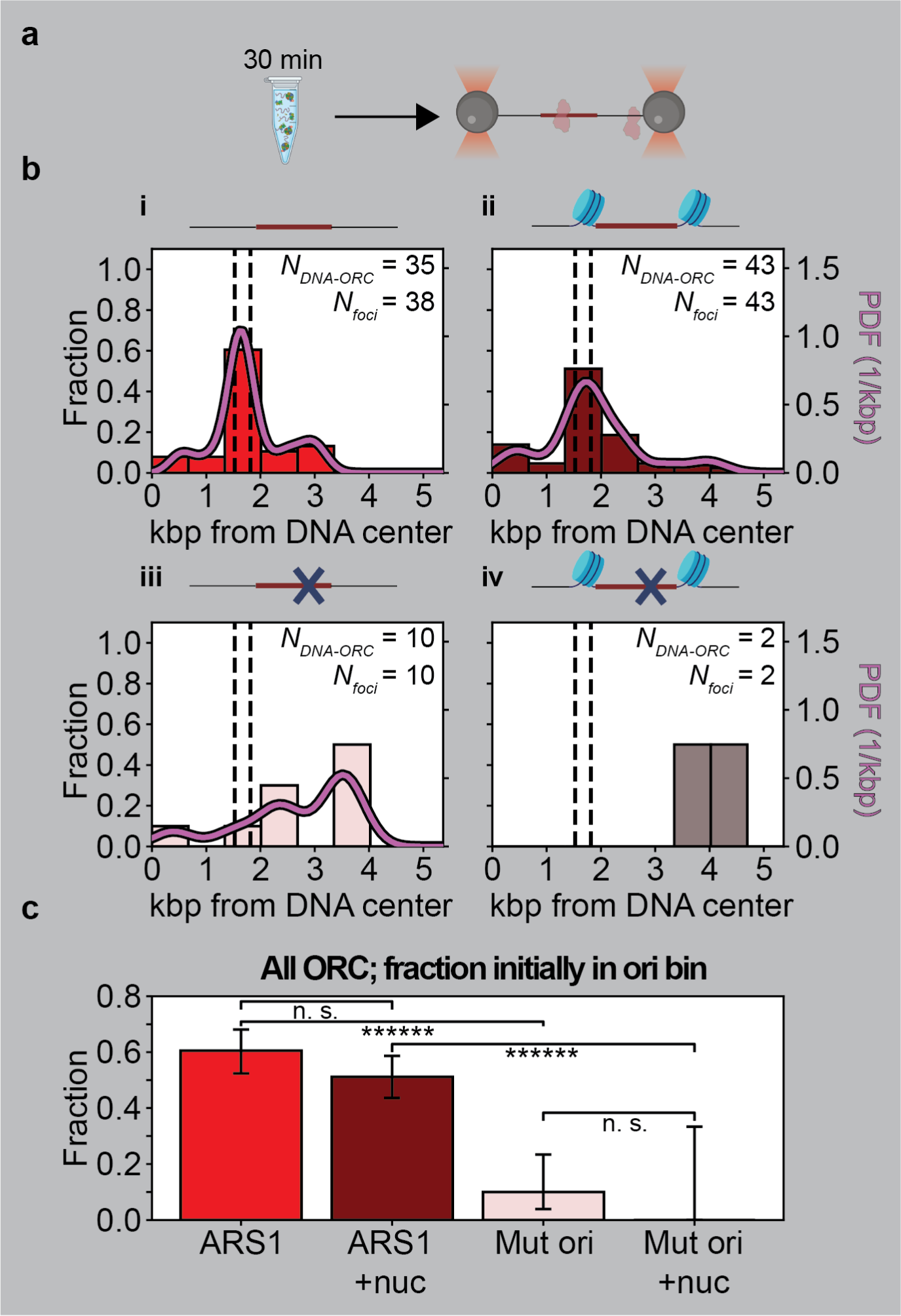
Spatial distribution of stably bound ORC on 10.4 kbp DNA containing a chromatinized origin. **(a)** Incubation method pertaining to data in panels (b-c). ORC and Cdc6 are incubated with DNA molecules (chromatinized or not, as indicated) in bulk for 30 min in the presence of ATP. Subsequently, the DNA-protein construct is flushed into the single-molecule flow cell, tethered, and moved into the buffer channel for confocal scanning. **(b)** (i) Spatial distribution of ORC on DNA containing a non-chromatinized origin as deduced from the diffraction-limited red spots (*N_foci_*) collected from 35 distinct DNA molecules (*N_DNA-ORC_*) analyzed and displayed as in panel (b-i). (ii) Spatial distribution of ORC on DNA containing a chromatinized origin as deduced from the diffraction-limited red spots (*N_foci_*) collected from 43 distinct DNA molecules (*N_DNA-ORC_*) analyzed and displayed as in panel (b-i). (iii) Spatial distribution of ORC on DNA containing a non-chromatinized mutated origin as deduced from the diffraction-limited red spots (*N_foci_*) collected from 10 distinct DNA molecules (*N_DNA-ORC_*) analyzed and displayed as in panel (b-i). (iv) Spatial distribution of ORC on DNA containing a chromatinized mutated origin as deduced from the red diffraction-limited spots (*N_DNA-ORC_*) collected from 2 distinct DNA molecules (*N_DNA-ORC_*) analyzed and displayed as in panel (b-i). The dashed lines indicate the location of the NPSs, and the solid curve indicates the kernel density estimation of the data. **(c)** ORC occupancy probability for the bins containing the (chromatinized or not) origin in panels (b-i)-(b-iv). Error bars are one-sigma Wilson confidence intervals. Statistical significance is determined by a two-sided binomial test. n.s. not significant, ****** *p* < 0.0000005.

Following this experimental approach, on bare DNA molecules containing the ARS1 origin we observed either DNA molecules devoid of ORC (*N_DNA-no ORC_* = 10, 21% of total), or DNA molecules (*N_DNA-ORC_* = 38, 79% of total) with ORC. Using the latter, we plotted the spatial distribution of ORC (**Figure 4b-i**). We found that 61% of the ORC foci were localized in the origin bin (*N_foci_* = 38, *N_foci in origin_* = 23), which emphasizes the preference of stable ORC binding at the origin compared to the other DNA sequences in the 10.4 kbp DNA^15^. Stoichiometric analysis showed no difference with the experiments reporting on rapid ORC binding (compare **Supplementary Figure 2b-i**, **Supplementary Figure 4.2b-i**). On DNA molecules with a chromatinized ARS1 origin, the overall fraction of DNA molecules containing ORC foci was reduced by a factor of two, to 41% (*N_DNA-no ORC_* = 74, *N_DNA-ORC_* = 52). However, we observed a similar fraction (51%) of ORC foci localized in the origin bin (*N_foci_* = 43, *N_foci in origin_* = 22, **Figure 4b-ii**), indicating that chromatinization of ARS1 did not enhance the likelihood of finding ORC stably bound in the origin bin relative to elsewhere on the DNA. Interestingly, stoichiometric analysis showed that such conditions led to an increase in the number of ORC molecules per focus (**Supplementary Figure 4.2b-ii**). When these experiments were repeated on DNA molecules with the mutated origin, we predominantly found DNA molecules devoid of ORC, irrespective of whether nucleosomes were present (*N_DNA-no ORC_* = 45, *N_DNA-ORC_* = 2, **Figure 4b-iv**) or not (*N_DNA-no ORC_* = 32, *N_DNA-ORC_* = 10, **Figure 4b-iii**). This suggested that stable ORC binding probed under these experimental conditions is predominantly associated with DNA sequence. For the four conditions probed, ORC binding in the bin containing the origin (chromatinized or not) was subjected to tests of statistical significance (**Figure 4c**). These results indicate that statistically significant enhancement of stable ORC binding depends on the presence of ARS1, but not on the presence of chromatin.

Previous studies have shown direct interactions between ORC and nucleosomes, suggesting a potential for nucleosome remodeling by ORC^11, 24^. Thus, we assessed whether in our experiments the presence of ORC and Cdc6 impacted the spatial distribution and stoichiometry of H2A. For reference, following a 30 min buffer-only bulk incubation of the DNA with a chromatinized ARS1 origin (*N_DNA_* = 51), 80% of H2A foci were found in the origin bin (*N_foci_* = 45, *N_foci in origin bin_* = 36), and the best fit of the H2A stoichiometry distribution yielded *p_h2a_* = 0.73 and *p_occupancy_* = 0.98 (**Supplementary Figure 4.1**), values similar to those obtained without such bulk incubation (**Figure 1c**). When the bulk incubation was performed in the presence of ORC and Cdc6 (*N_DNA_* = 126), however, the fraction of H2A foci in the origin bin was reduced to 49% (*N_foci_* = 134, *N_foci in origin bin_* = 66) (**Supplementary Figure 4.3b**), and the best fit of the H2A stoichiometry distribution yielded *p_h2a_* = 0.81 and *p_occupancy_* =0.52. The latter parameters suggested a maintenance of H2A-H2B dimer stability but a less complete NPS occupancy. When the bulk incubation in the presence of ORC and Cdc6 was performed on DNA molecules with a chromatinized mutated origin (*N_DNA_* = 47), an intermediate value of 64% of H2A foci were found in the origin bin (*N_foci_* = 56, *N_foci in origin bin_* = 36) (**Supplementary Figure 4.3c**), and the best fit of the H2A stoichiometry distributions of H2A yielded *p_h2a_* = 0.60 and *p_occupancy_* = 1.00. The latter parameters indicated a reduction in H2A-H2B dimer stability without a change in NPS site occupancy. In summary, these results show that under these more extended incubation conditions, the presence of ORC and Cdc6 can influence nucleosome positioning and stability.

### Nucleosomes flanking the origin permit MCM recruitment but restrict subsequent motion

Having probed both rapid and stable ORC binding and established that the presence of ARS1 was beneficial to both but the presence of nucleosomes only to the former, we wanted to test how the presence of nucleosomes impacted the loading of MCM. We therefore set out to probe the spatial positioning, stoichiometry, and dynamics of MCM on a chromatinized origin. To visualize MCM, we used JF646-labeled MCM in loading reactions. MCM was labeled by introducing a HaloTag on the N-terminus of its Mcm3 subunit, and JF646-MCM performed normally in a bulk loading assay (**Supplementary Figure 1.1**). We then incubated ORC, Cdc6 and MCM/Cdt1 with DNA in bulk, without tethering to beads, for 30 min^5, 15^ as described above for the experiments that probed for stable ORC binding. We next introduced these pre-incubated DNA molecules into the flow cell, and imaged JF646-MCM as described above (**Figure 5a**).

**Figure 5.**
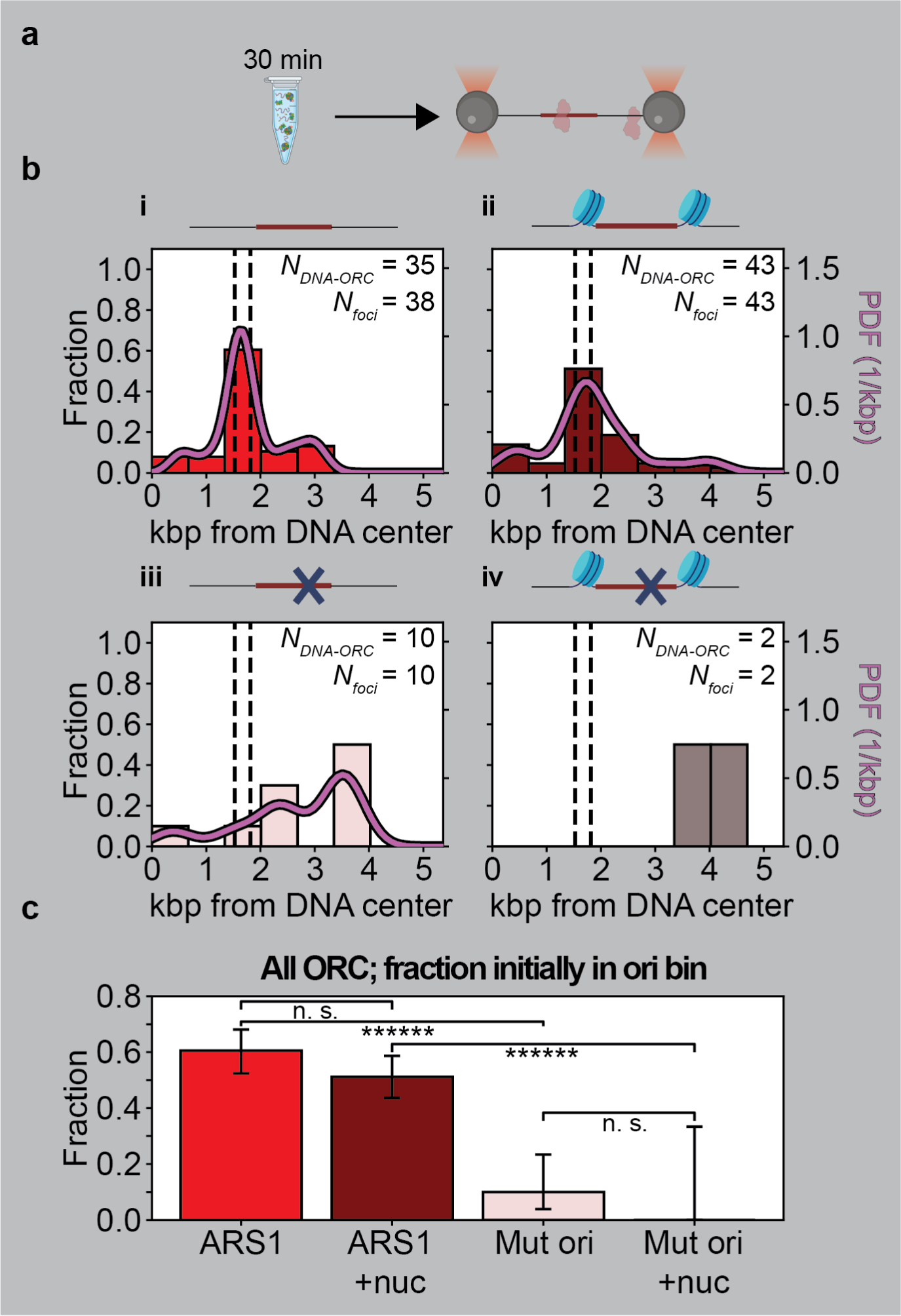
Nucleosomes spatially constrain MCM at the origin. **(a)** ORC, Cdc6, and Mcm2-7/Cdt-1 are incubated with 10.4 kb DNA molecules (chromatinized or not, as indicated) in bulk for 30 min in the presence of ATP. Subsequently, the DNA-protein construct is flushed into the single-molecule flow cell, tethered and moved into the buffer channel for confocal scanning with an acquisition frequency of one frame every 0.01 min (*t*_f_ = 0.01 min). **(b)** (i) Spatial distribution of Mcm2-7 on DNA containing a non-chromatinized origin as deduced from the diffraction-limited red spots (*N_foci_*) collected from 44 distinct DNA (*N_DNA-MCM_*). The dashed lines indicate the location of the NPSs, and the solid curve indicates the kernel density estimation of the data. (ii) Spatial distribution of Mcm2-7 on DNA containing a chromatinized origin as deduced from the diffraction-limited red spots (*N_foci_*) collected from 77 distinct DNA molecules (*N_DNA-MCM_*) analysed and displayed as in panel (b-i). (iii) Spatial distribution of Mcm2-7 on DNA containing a non-chromatinized mutated origin as deduced from the diffraction-limited red spots (*N_foci_*) collected from 65 distinct DNA molecules (*N_DNA-MCM_*) analyzed and displayed as in panel (b-i). (iv) Spatial distribution of Mcm2-7 on DNA containing a chromatinized mutated origin as deduced from the diffraction-limited red spots (*N_foci_*) collected from 71 distinct DNA molecules (*N_DNA-MCM_*) analyzed and displayed as in panel (b-i). **(c)** Mcm2-7 occupancy probability for the bins containing the (chromatinized or not) origin for the datasets in panels (b-i)-(b-iv). Error bars are one-sigma Wilson confidence intervals. Statistical significance is determined by a two-sided binomial test: n.s. not significant and ****** *p* < 0.0000005. **(d)** ORC, Cdc6, and Mcm2-7/Cdt-1 are incubated with 10.4 kb chromatinized DNA molecules in bulk for 30 min in the presence of ATP. Subsequently, the DNA-protein construct is flushed into the single-molecule flow cell, tethered and moved into the buffer channel for confocal scanning with an acquisition frequency of one frame per minute (*t*_f_ = 1 min). **(e)** Histograms of the relative displacements (Δ*x* = 2 min) between 2 min intervals (Δ*t* = 2 min) for (i) dCas9 molecules over the entire lifetime of the fluorophore for dCas9 molecules, (ii) for Mcm2-7 on DNA containing a non-chromatinized origin or (iii) chromatinized origin. For all three panels, acquisition is continued until the fluorophore is bleached (∼20 min). *N_foci_*, number of molecules; *N*_Δ_*_x_*, number of relative displacements; *σ*, standard deviation.

In the absence of nucleosomes, MCM foci were broadly distributed about the ARS1 origin (**Figure 5b-i**), as previously described^15^. Chromatinization of ARS1 origin, however, resulted in a strikingly different situation: MCM foci were now mainly localized in the origin bin (**Figure 5b-ii**). Repeating this experiment on bare DNA containing the mutated origin again resulted in a broad distribution of MCM (**Figure 5b-iii**), this time with a slight peak in the same bin as was observed in the experiments that probed rapid ORC binding (**Figure 2b-iii**). Chromatinization of this mutated origin did not significantly increase the presence of MCM in the origin bin (**Figure 5b-iv**). For the four types of DNA molecules probed, MCM binding in the bin containing the origin (chromatinized or not) was subjected to tests of statistical significance (**Figure 5c**). These results indicate that chromatinization of the ARS1 origin, but not of a DNA segment without ARS1, statistically significantly increases the population of MCM bound there.

We also examined the stoichiometry of MCM foci bound to four different types of DNA molecules tested under these four conditions. Stoichiometric analysis showed that the MCM foci predominantly contained 1 or 2 molecules for all cases (**Supplementary Figure. 5.1**). However, foci found on DNA molecules that contained the chromatinized ARS1 origin exhibited the highest fraction of foci in the origin bin that contained 2 MCM molecules (53%, right plot in **Supplementary Figure 5.1-ii**). We discuss this further below, together with the overall observation that the spatial distributions of MCM resembled those obtained for rapid ORC binding (**Figure 2**) more closely than those obtained for stable ORC binding (**Figure 4**).

As our experiments showed that the presence of ORC and Cdc6 during 30 min incubations in bulk impacted nucleosome positioning and stability (**Supplementary Figure 4.3**), we also monitored nucleosome positioning and stability upon the inclusion of MCM. On DNA molecules with a chromatinized ARS1 (*N_DNA_* = 110), 83% of H2A foci were found in the origin bin (*N_foci_* = 81, *N_foci in origin bin_* = 67), and the best fit of the H2A stoichiometry distribution yielded *p_h2a_* = 0.84 and *p_occupancy_* = 0.89 (**Supplementary Figure 5.2b**), which are all values similar to those obtained following incubation with buffer alone. However, on DNA with a chromatinized mutated origin (*N_DNA_* = 123), only 50% of H2A foci were located in the origin bin (*N_foci_* = 70, *N_foci in origin bin_* = 140), and the best fit of the H2A stoichiometry distribution yielded *p_h2a_* = 0.91 and *p_occupancy_* = 0.40. The latter parameters suggested a maintenance of H2A-H2B dimer stability but a less complete NPS site occupancy. We speculate that increased MCM loading on a chromatinized ARS1 origin (**Supplementary Figure 5.2b**) relative to the chromatinized mutated origin (**Supplementary Figure 5.2c**) limits the continued access of ORC and Cdc6 to nucleosomes, and hence their impact (see also Discussion).

Lastly, we tested whether a pair of nucleosomes surrounding ARS1 can prevent MCM, once loaded, from diffusing outwards away from ARS1. To test this directly, we monitored MCM position over an extensive duration of time by repeated the preceding experiments but imaging JF646-MCM at a 100-fold slower rate (one frame per min, **Figure 5d**). This allowed us to probe for MCM motion occurring on timescales up to approximately 15 min. We measured the relative displacements of JF646-MCM foci every 2 min (or 2 frames) until JF646 was no longer visible due to photobleaching (lifetime of the JF646 dye on DNA-bound dCas9 under identical imaging conditions was 1.23 x 10^3^ s). These data were benchmarked relative to the displacement distribution of dCas9 (**Figure 5e-i**) obtained under identical experimental conditions, which was centered about 0 and had a width of *σ* = 0.061 kbp that derived from experimental noise. The distribution of displacements by MCM on DNA molecules with a bare ARS1 had a substantially increased width relative to dCas9 (**Figure 5f-ii**; *σ* = 0.398 kbp), indicating one-dimensional diffusion of the MCM foci as previously observed^15^. However, on DNA containing the chromatinized ARS1, the distribution of displacements by MCM again had a narrow width (**Figure 4f-iii**; *σ* = 0.153 kbp). This indicated that one-dimensional diffusion of MCM away from the ARS1 origin of replication was hampered by the presence of flanking nucleosomes.

## DISCUSSION

Chromatin replication starts from origins that are flanked by nucleosomes. We have studied at the single-molecule level to understand how the presence of nucleosomes impacts the binding and mobility of ORC in the vicinity of the origin and subsequently, the recruitment and mobility of MCM.

### Nucleosomes facilitate rapid binding of ORC to the origin and decrease its mobility

Our experiments first probed the rapid binding of yeast ORC to DNA following a short incubation period in the single-molecule instrument (**Figure 2**) and the subsequent motion of these molecules (**Figure 3**). As our previous work on bare DNA has shown, mobile ORC molecules can contribute to origin recognition by scanning the DNA – given the measured diffusion constant of the fast ORC population (**Figure 3b-i**), even during a short 5 s incubation period an individual ORC molecule can scan ∼1.2 kbp on bare DNA, which contributes to its observed binding at many different sites along the DNA (**Figure 2b-i**). Repeating these experiments in the presence of a chromatinized origin showed that the presence of nucleosomes provides a mild increase in local ORC binding (**Figure 2b-ii, c**) and a reduction in ORC mobility (**Figure 3b-ii, c**). The latter can derive from spatial confinement of ORC by nucleosomes, direct interactions between ORC and nucleosomes, or a combination thereof. A reduction in ORC mobility can furthermore account for the observed increase in local ORC binding: for example, if nucleosomes similarly reduced the influx of mobile ORC molecules into the origin region and their efflux out of it, then direct interactions between ORC and nucleosomes could account for the observed increase in local ORC binding. Certainly, our data suggest that in the yeast system ORC molecules can readily locate a chromatinized origin via binding from solution even if their one-dimensional diffusion is blocked; whether the same holds true for higher eukaryotes that lack sequence-specific origins remains to be determined.

### Stable binding of ORC is primarily enabled by the origin of replication

We next asked whether nucleosomes affected the stable binding of ORC to DNA following bulk incubation of ORC with DNA molecules over an extended duration. Examination in the single-molecule instrument revealed that a bare ARS1 origin alone sufficed to retain stably bind ORC molecules to DNA (**Figure 4a**); no further increases in the yield of stable ORC were observed with a chromatinized ARS1 origin (**Figure 4b**). Interestingly, while DNA molecules that contained a mutated origin could recruit rapidly bound ORC (**Figure 2d,e**), they could not stably retain ORC, irrespective of whether nucleosomes surrounded the origin (**Figure 4d,e**). This most likely suggests that rapidly but nonspecifically recruited ORC molecules unbind from the DNA directly into solution^25^, as our direct tracking of the motion of ORC does not provide experimental evidence that ORC can bypass flanking nucleosomes (**Figure 3**). These experiments thus showed that ARS1 is required for stable ORC binding^15^ even in the presence of nucleosomes. This contrast with results from another study^26^ that suggested that a specific sequence might not be necessary and found ORC bound specifically to nucleosomes. Such discrepancies may have resulted from differences in the experimental preparations employed. Nevertheless, our findings are in line with previous findings that show that origin sequences are required for yeast replication *in vitro* on chromatinized DNA and *in vivo* in budding yeast^27, 28^.

Stoichiometric analysis of either rapidly or stably bound ORC molecules bound to the region containing the ARS1 origin indicated an increase in ORC stoichiometry upon chromatinization (**Supplementary Figure 2**). We computed the resulting ratio of 1 ORC molecule:2 ORC molecules to be ∼1:3, very similar to the ratio previously determined using cryo-EM^16^. An increase in ORC stoichiometry upon chromatinization of ARS1 could result from a similar influx of ORC to the origin but a reduced efflux as the diffusion of ORC molecules away from the origin^15^ is reduced by the presence of nucleosomes, or an increased influx of ORC to the origin due to direct interactions between ORC and nucleosomes^26, 29^, or a combination of these effects.

### Influence of ORC and MCM on nucleosome remodeling at the origin of replication

Previous studies have indicated that ORC has properties of a chromatin remodeler^12–14, 24^, either due to direct interaction of ORC with the nucleosomes or due to DNA bending at the origin^30^. Indeed, we confirmed that extended incubation of chromatinized substrates with ORC and Cdc6 could broaden the spatial distribution and/or reduce the stoichiometry of H2A histones embedded within our nucleosomes (**Supplementary Figure 4.3**), in agreement with published work reporting in vitro remodeling of H2A-H2B dimers by ORC^24^. Interestingly, our data monitoring the position and stoichiometry distributions of H2A following an extended incubation with all loading factors, however, show no changes therein on DNA containing a chromatinized ARS1 origin (**Supplementary Figure 5.2**). This suggests a protective role for MCM in the maintenance of epigenetic chromatin states prior to the firing of DNA replication^31^.

### Loading of MCM onto a chromatinized origin

Our results showed that the lack of a replication origin inhibits the stable binding of ORC to DNA (**Figure 4**), but not the rapid binding of ORC (**Figure 2**) or the loading of MCM (**Figure 5**). This implies that stable binding of ORC at ARS1 is not a prerequisite to MCM loading, which is consistent with biochemical experiments showing that MCM loading can occur independently of specific origins^5^. Furthermore, our measurements of the spatial distributions of MCM indicate that for the four types of DNA molecules tested, there is a probability of finding MCM molecules all along the DNA molecule. Several factors underlie this behavior. First, it is the one just mentioned: ORC binding is not strictly sequence-specific even in yeast, and thus MCM loading can occur at different DNA sequences. Indeed, the spatial distribution of MCM molecules (**Figure 5**) showed similar features to the spatial distributions of rapid ORC binding (**Figure 2**). Second, both bulk biochemical assays^4, 5^ and our previous single-molecule studies^15^ have shown that loaded MCM can undergo linear diffusion on DNA. In the buffer conditions employed, the diffusion constant of MCM equals 0.0008 ± 0.0002 kbp^2^ s^-5^ (**Figure 5e-ii**), from which we estimate that diffusion of MCM during the extended incubation period contributed ∼1.4 kbp to the broadening of the observed spatial distribution.

On DNA molecules containing a chromatinized ARS1, we found that the probability of finding MCM proteins in the vicinity of the origin was increased by more than two-fold compared to that on DNA molecules containing bare ARS1. Given that this increase is larger for MCM (**Figure 5c**) than for rapidly bound ORC (**Figure 2c**), this data alone could suggest that the presence of nucleosomes prevents the diffusion of MCM molecules away from the origin. The lack of a similar increase in the probability of MCM foci close to the origin due to the chromatinization of a mutated origin, compare **Figure 4e** to **Figure 4d** (despite a corresponding mild increase in rapid ORC binding, compare **Figure 2e** to **Figure 2d**) is consistent with previous work showing that nucleosomes reduced non-specific loading of MCM on a mutated origin^14, 32^.

Stoichiometric analysis reveals that the detected MCM foci in region of the ARS1 origin predominantly have a stoichiometry of 1 or 2. In the presence of the nucleosomes, however, we clearly observed an increased fraction of foci with a stoichiometry of 2 (**Supplementary Figure 5.1b**). While in our experimental configuration a measured MCM stoichiometry of 2 within a diffraction-limited focus cannot distinguish an MCM double hexamer from two single MCM hexamers, our results are consistent with previous work showing that a chromatinized ARS1 origin favors the formation of the MCM/ORC (MO) intermediate complex and, consequently, the loading of MCM double hexamers^16^.

### MCM spatially constrained to a chromatinized origin on long timescales

Our experimental tracking of MCM position over long incubation times shows that the presence of nucleosomes spatially constrains MCM (**Figure 5e**). Does this spatial confinement help to explain how the presence of origin-flanking nucleosomes favors pre-RC formation^11, 12^? Such spatial confinement will limit the diffusion of MCM single hexamers^15, 23^ and thereby reduce the likelihood of MCM double hexamer formation through the encounter of properly oriented, diffusing MCM single hexamers^23^. Possibly, this favors MCM double hexamer formation via the MO intermediate^16^. Future investigations will further investigate the confinement of MCM by nucleosomes, as it could potentially contribute to the recycling of licensing factors in cases of DNA replication stress, where reloading of factors is not possible. replisome assembly at origins of replication flanked by nucleosomes.

## CONCLUSIONS

Our findings provide new insights into the role of chromatin in DNA replication initiation. We demonstrate that the presence of nucleosomes surrounding origins contributes to the loading and subsequent spatial constraint of MCM hexamers there, which likely favors subsequent CMG formation and the initiation of DNA replication at replication origins. Future investigation will focus on the quantification of these downstream events. This research will additionally open up new avenues to further explore the role of chromatin in origin firing and the mechanisms of chromatin replication.

## METHODS

### Biological Materials

#### Protein purification and labeling

##### Histone octamers

Plasmids pCDFduet.H2A-H2B and pET-Duet.H3-H4, expressing *S. cerevisiae* histones H2A, H2B, H3 and H4 were kindly provided by Dr. Martin Singleton (Francis Crick Institute, London). Lysine 120 of H2A was changed to cysteine by site-directed mutagenesis to facilitate fluorescent labeling. These histones were co-expressed in *Escherichia coli* strain BL21-codonplus-DE3-RIL (Agilent), and the histone octamer with the K120>C mutation in histone H2A was purified according to Ref^33^. Cells were grown to a density with an OD^600^ of 0.3-0.5, and expression of the histones was induced with 400 µM isopropyl 1-thio-ß-D-galactopyranoside (Santa Cruz Biotechnology Inc) for 16 h at 17°C while shaking at 180 rpm. Cells were lysed by sonication in a Qsonica Q500 sonicator for 2 min with cycles of 5 s on and 5 s off and an amplitude of 40%, in histone lysis buffer (0.5 M NaCl, 20 mM Tris-HCl pH 8, 0.1 mM EDTA, 1 mM DTT, 0.3 mM PMSF, and protease cOmplete inhibitor). Supernatant containing the histone octamers was purified on a 5 mL Hi-Trap Heparin column (Cytiva) and eluted with a gradient of 0.5–2 M NaCl in 20 mM Tris-HCl pH 8, 0.1 mM EDTA, 1 mM DTT. Peak fractions were analyzed on a 12% SDS-PAGE, and octamer-containing fractions were further purified on a Superdex 200 increase (Cytiva) using histone GF buffer (2 M NaCl, 20 mM Tris-HCl pH 8, 0.1 mM EDTA, and 1 mM DTT). Peak fractions were analyzed on a 12% SDS-PAGE gel, and fractions containing histone octamers were pooled and concentrated in an Amicon Ultra-4 Ultracell 30kDa centrifugal filter (Merck-Millipore #UFC803024). The protein concentration was determined with Bio-Rad Protein Assay Dye Reagent (Bio-Rad # 5000006).

##### Cdc6

*S. cerevisiae* Cdc6 protein expression was induced in BL21-CodonPlus(DE3)-RIL cells (Agilent #230245) transformed with pGEX-6P-1 wt GST-cdc6 using 400 µM IPTG for 16 h at 16 °C. Cells were harvested in Cdc6 lysis buffer (50 mM K_X_PO_4_ pH 7.6, 150 mM KOAc, 5 mM MgCl_2_, 1% Triton X-100, 2 mM ATP, cOmplete^TM^ EDTA-free Protease Inhibitors (Sigma-Aldrich #5056489001), and 1 mM DTT) and sonicated in a Qsonica Q500 sonicator for 2 min with cycles of 5 s and 5 s off and an amplitude of 40%. After centrifugation, Cdc6 protein was purified from the supernatant by incubating for 1 h at 4 °C with glutathione beads Fastflow (GE Healthcare #17-5132-02). The beads were washed 20 times with 5 ml Cdc6 lysis buffer, and Cdc6 was released from the beads by digestion with Precision protease (GE Healthcare #27-0843-01) at 4 °C for 16 h. Subsequently, the Cdc6 eluate was diluted with Cdc6 dilution buffer (50 mM K_X_PO_4_ pH 7.6, 5 mM MgCl_2_, 0.1% Triton X-100, 2 mM ATP, and 1 mM DTT) to a final KOAc concentration of 75 mM and incubated with hydroxyapatite Bio gel HTP (Bio-Rad #130-0402) for 45 min at 4 °C. The beads were washed five times with Cdc6 wash buffer (50 mM K_X_PO_4_ pH 7.6, 75 mM KOAc, 5 mM MgCl_2_, 0.1% Triton X-100, 2 mM ATP, and 1 mM DTT), and then washed five times with Cdc6 rinse buffer (50 mM K_X_PO_4_ pH 7.6, 150 mM KOAc, 5 mM MgCl_2_, 15% glycerol, 0.1% Triton X-100, and 1 mM DTT). Next, Cdc6 was eluted from the column in 1-ml fractions with Cdc6 elution buffer (50 mM KXPO4 pH 7.6, 400 mM KOAc, 5 mM MgCl_2_, 15% glycerol, 0.1% Triton X-100, and 1 mM DTT). Finally, fractions containing Cdc6 were pooled, dialyzed twice for 1 h against Cdc6 dialysis buffer (25 mM HEPES-KOH pH 7.6, 100 mM KOAc, 10 mM MgOAc, 10% glycerol, and 0.02% NP40 substitute) in a 10 kDa cut off Slide-A-Lyzer Cassette (Thermo Scientific #66380), and concentrated in an Amicon Ultra-4 Ultracell 30kDa centrifugal filter (Merck-Millipore #UFC803024). Aliquots were snap frozen and stored at −80 °C. The protein concentration was determined with Bio-Rad Protein Assay Dye Reagent (Bio-rad # 5000006).

##### ORC and Halo-tagged ORC

ORC complex with a CBP-TEV tag on orc1 was purified from *S. cerevisiae* strain ySDORC, and ORC complex with a CBP-TEV-Halo tag on orc3 was purified from strain yTL158. Cells were seeded at a density of 2*10^7^ cells per ml in YP medium (1% yeast extract and 2% peptone), supplemented with 2% raffinose and grown at 30 °C and 180 rpm until a density of 3-5*10^7^ cells/ml was reached. Then cells were arrested in G1 by adding 100 ng/ml α-mating factor (Tebu-Bio #089AS-60221-5) for 3 h followed by the addition of 2% galactose for 3 h to induce the expression of ORC. Cells were harvested by centrifugation and washed with ORC lysis buffer (25 mM HEPES-KOH pH 7.6, 0.05% NP-40 substitute, 10% glycerol, 0.1 M KCl, and 1 mM DTT). After centrifugation, cells were suspended in ORC lysis buffer supplemented with protease inhibitors (cOmplete^TM^ EDTA-free Protease Inhibitors (Sigma-Aldrich #5056489001) and 0.3 mM PMSF) and dropped into liquid nitrogen. The frozen droplets were ground in a freezer mill (6875 SPEX) for 6 cycles (run time 2 min and cool time 1 min with a rate of 15 cps), and the resulting powder was suspended in ORC lysis buffer supplemented with protease inhibitors. The lysate was cleared in a Beckman-Coulter ultracentrifuge (type Optima L90K with rotor TI45) for 1 h at 235.000 g at 4 °C. The cleared lysate was supplemented with CaCl_2_ to a final concentration of 2 mM and with KCl to a final concentration of 0.3 M and then incubated for 1 h at 4 °C with washed Sepharose 4B Calmodulin beads (GE Healthcare #17-0529-01) in a spinning rotor. The beads were washed 20 times with 5 ml ORC binding buffer (25 mM HEPES-KOH pH 7.6, 0.05% NP-40 substitute, 10% glycerol, 0.3 M KCl, 2 mM CaCl_2_, and 1 mM DTT), and the protein complex was eluted from the beads with ORC elution buffer (25 mM HEPES-KOH pH 7.6, 0.05% NP-40 substitute, 10% glycerol, 0.3 M KCl, 2 mM EDTA, 2 mM EGTA, and 1 mM DTT). ORC-containing fractions were pooled, concentrated in an Amicon Ultra-4 Ultracell 30kDa centrifugal filter (Merck-Millipore #UFC803024), and applied to a Superose 6 increase 10/300 GL column (GE Healthcare #29-0915-96) equilibrated in ORC GF buffer (25 mM HEPES-KOH pH 7.6, 0.05% NP-40 substitute, 10% glycerol, 0.15 M KCl, and 1 mM DTT). Peak fractions were pooled and concentrated in an Amicon Ultra-4 Ultracell 30kDa centrifugal filter (Merck-Millipore #UFC803024). Aliquots were snap frozen and stored at −80°C. The protein concentration was determined with Bio-Rad Protein Assay Dye Reagent (Bio-Rad # 5000006).

##### Mcm2-7/Cdt1 and Halo-tagged Mcm2-7/Cdt1

Mcm2-7/Cdt1 complex with a CBP-TEV tag on mcm3 was purified from *S. cerevisiae* strain yAM33, and Mcm2-7/Cdt1 complex with a CBP-TEV-Halo tag on mcm3 was purified from strain yTL001. Cells were grown, and Mcm2-7/Cdt1 expression was induced as described for ORC. Cells were harvested by centrifugation and washed with MCM lysis buffer (45 mM HEPES-KOH pH 7.6, 0.02% NP-40 substitute, 10% glycerol, 100 mM KOAc, 5 mM MgOAc, and 1 mM DTT). After centrifugation, cells were suspended in MCM lysis buffer supplemented with protease inhibitors (cOmplete^TM^ EDTA-free Protease Inhibitors (Sigma-Aldrich #5056489001) and 0.3 mM PMSF) and dropped into liquid nitrogen. The frozen droplets were ground in a freezer mill (6875 SPEX) for 6 cycles (run time 2 min and cool time 1 min at a rate of 15 cps), and the resulting powder was suspended in MCM lysis buffer supplemented with protease inhibitors. The lysate was cleared in a Beckman-Coulter ultracentrifuge (type Optima L90K with rotor TI45) for 1 h at 235.000 g and 4 °C. The cleared lysate was supplemented with CaCl_2_ to a final concentration of 2 mM and then incubated for 1 h at 4 °C with washed Sepharose 4B Calmodulin beads (GE Healthcare #17-0529-01) in a spinning rotor. The beads were washed 20 times with 5 ml MCM binding buffer (45 mM HEPES-KOH pH 7.6, 0.02% NP-40 substitute, 10% glycerol, 100 mM KOAc, 5 mM MgOAc, 2 mM CaCl_2_, and 1 mM DTT), and the protein complex was eluted from the beads with MCM elution buffer (45 mM HEPES-KOH pH 7.6, 0.02% NP-40 substitute, 10% glycerol, 100 mM KOAc, 5 mM MgOAc, 1 mM EDTA, 2 mM EGTA, and 1 mM DTT). Mcm2-7/Cdt1-containing fractions were pooled, concentrated in an Amicon Ultra-4 Ultracell 30kDa centrifugal filter (Merck-Millipore #UFC803024), and applied to a Superose 6 increase 10/300 GL column (GE Healthcare #29-0915-96) equilibrated in MCM GF buffer (45 mM HEPES-KOH pH 7.6, 0.02% NP-40 substitute, 10% glycerol, 100 mM KOAc, 5 mM MgOAc, and 1 mM DTT). Peak fractions were pooled and concentrated in an Amicon Ultra-4 Ultracell 30kDa centrifugal filter (Merck-Millipore #UFC803024). Aliquots were snap frozen and stored at −80 °C. Protein concentration was determined with Bio-Rad Protein Assay Dye Reagent (Bio-Rad # 5000006).

##### dCas9-Halo

Halo-tagged dCas9 protein expression was induced in BL21-CodonPlus(DE3)-RIL cells (Agilent #230245) transformed with pET302-6His-dCas9-halo (Addgene #72269) using 400 µM IPTG for 16 h at 16 °C. Cells were harvested in dCas9 lysis buffer (50 mM Na_x_PO_4_ pH 7.0, 300 mM NaCl and protease inhibitors (cOmplete^TM^ EDTA-free Protease Inhibitors (Sigma-Aldrich #5056489001) plus 0.3 mM PMSF)) and sonicated in an Qsonica Q500 sonicator for 2 min with cycles of 5 s on and 5 s off and an amplitude of 40%. After centrifugation, dCas9-Halo protein was purified from the supernatant by incubating for 2 h at 4 °C with Ni-NTA agarose (Qiagen #30210). The beads were washed 10 times with 5 ml dCas9 wash buffer I (50 mM Na_x_PO_4_ pH 7.0 and 300 mM NaCl) and 3 times with dCas9 wash buffer II (50 mM Na_x_PO4 pH 7.0, 300 mM NaCl, and 20 mM Imidazole pH 7.6), and dCas9-Halo was eluted from the agarose beads with dCas9 elution buffer (50 mM Na_x_PO_4_ pH 7.0, 300 mM NaCl, and 150 mM Imidazole pH 7.6). Subsequently, dCas9-Halo eluate was dialyzed twice for 1 h against dCas9-dialysis buffer (50 mM HEPES-KOH pH 7.6, 100 mM KCl, and 1 mM DTT) in a 10kDa cut off Slide-A-Lyzer Cassette (Thermo Scientific #66380) and applied to a Hi Trap SP HP column (GE Healthcare #17-1151-01) equilibrated with dCas9 dialysis buffer. The dCas9-Halo protein was eluted from the column with dialysis buffer with a KCl gradient ranging from 100 mM up to 1 M. The dCas9-Halo-containing fractions were pooled, concentrated in an Amicon Ultra-4 Ultracell 30kDa centrifugal filter (Merck-Millipore #UFC803024), and applied to a Superdex 200 increase 10/300 GL column (GE Healthcare #28-9909-44) equilibrated in cas9 GF buffer (50 mM HEPES-KOH pH 7.6, 150 mM KCl, and 1 mM DTT). Peak fractions were pooled and concentrated in an Amicon Ultra-4 Ultracell 30kDa centrifugal filter (Merck-Millipore #UFC803024). Aliquots were snap frozen and stored at −80 °C. The protein concentration was determined with Bio-Rad Protein Assay Dye Reagent (Bio-Rad # 5000006).

##### dCas9-Cys

dCas9-Cys protein expression was induced in BL21-CodonPlus(DE3)-RIL cells (Agilent #230245) transformed with plasmid 10xHis-MBP-TEV-S. pyogenes dCas9 M1C D10A C80S H840A C574S (Addgene #60815) with 400 µM IPTG for 5 h at 20 °C. Cells were harvested in dCas9-cys lysis buffer (20 mM Tris-HCl pH 8.0, 500 mM NaCl, 1mM DTT, and protease inhibitors (cOmpleteTM EDTA-free Protease Inhibitors (Sigma-Aldrich #5056489001) plus 0.3 mM PMSF)) and sonicated in an Qsonica Q500 sonicator for 2 min with cycles of 5 s on and 5 s off and an amplitude of 40%. After centrifugation, dCas9-Cys protein was purified from the supernatant by incubating for 2 h at 4 °C with Ni-NTA agarose (Qiagen #30210). The beads were washed 10 times with 5 ml dCas9-Cys wash buffer (20 mM Tris-HCl pH 8.0, 250 mM NaCl, 20mM imidazole pH 7.6), and dCas9-Cys was eluted from the agarose beads with dCas9-Cys elution buffer (20 mM Tris-HCl pH 8.0, 250 mM NaCl, 150 mM imidazole pH 7.6). Subsequently, dCas9-Cys eluate was dialyzed twice for 1 h against dCas9-Cys-dialysis buffer (20 mM HEPES-KOH pH 7.6, 150 mM KCl, and 1 mM DTT) in a 10 kDa cut off Slide-A-Lyzer Cassette (Thermo Scientific #66380) and applied to a Hi Trap SP HP column (GE Healthcare #17-1151-01) equilibrated with dCas9-Cys dialysis buffer. The dCas9-Cys protein was eluted from the column with dCas9-Cys dialysis buffer with a KCl gradient ranging from 100 mM up to 1 M. The dCas9-Cys-containing fractions were pooled, concentrated in an Amicon Ultra-4 Ultracell 30kDa centrifugal filter (Merck-Millipore #UFC803024), and applied to a Superdex 200 increase 10/300 GL column (GE Healthcare #28-9909-44) equilibrated in cas9-Cys GF buffer (20 mM HEPES-KOH pH 7.6, 150 mM KCl, and 1 mM DTT). Peak fractions were pooled and concentrated in an Amicon Ultra-4 Ultracell 30 kDa centrifugal filter (Merck-Millipore #UFC803024). Aliquots were snap frozen and stored at −80 °C. The protein concentration was determined with Bio-Rad Protein Assay Dye Reagent (Bio-Rad # 5000006).

#### Protein labeling

Strains: To create Halo-tagged mcm3, the StuI and XmaI restriction sites in plasmid pENTR4-HaloTag (Addgene #W876-1) were changed into a silent mutation following standard cloning techniques using primers TL-019-TL-020 and TL-023-TL-024. The sequence was verified by sequencing using primers TL-021-TL-022. Then the HaloTag fragment was amplified from the mutated pENTR4-HaloTag by PCR with primers TL-025 and TL-026, which were extended with an XmaI site. This amplified HaloTag was digested with XmaI, gel-purified, and ligated into plasmid pRS306 CBP-TEV-mcm3-gal1-10 mcm2, which was digested with SgrAI and dephosphorylated with CIP, resulting in plasmid pRS306 CBP-TEV-mhalo-mcm3-gal1-10 mcm2. Proper integration of the HaloTag was confirmed by sequencing with primers (see Supplementary Table 1) TL-001, TL-002, TL-027, and TL-028. Yeast strain yTL001, which expresses MCM with a Halo-tagged mcm3, was created by linearizing plasmid pRS306 CBP-TEV-mhalo-mcm3-gal1-10-mcm2 with StuI and transforming it into yeast strain yJF21, which expresses Mcm4-7 and Cdt1 upon induction with galactose.

To create an ORC complex with a halo-tagged orc3, the CBP-TEV sites was removed from plasmid pRS306 orc1-gal1-10-orc2 through Gibson assembly (NEB #E2611L) using primers TL-441, TL-443, and TL-447. The sequence for the coding region of orc1 and orc2 was confirmed by sequencing using primers TL-084, TL-087, TL-119, and TL-136. Yeast strain yTL151, which expresses orc1, 2, 5, and 6 from a galactose-inducible promoter, was created by linearizing plasmid pRS306 orcl-gal1-10-orc2 v2 delta CBP-TEV with StuI and transforming it into yeast strain yTL070, which contains an inducible expression plasmid for orc5 and orc6.

Plasmid pRS303 CBP-TEV-halo-orc3 gal1-10 orc4 was generated by cloning the CBP-TEV-halo sequence from plasmid pRS306-CBP-TEV-halo-Pri1-Gal1-10 Pri2 into plasmid pRS303-orc3-Gal1-10 orc4 through Gibson assembly (NEB #E2611L) using primers TL-446, TL-447, TL-472, and TL-473). The sequence of CBP-TEV-halo-orc3 and orc4 was verified by sequencing using primers TL-063, TL-064, TL-449, and TL-470. Yeast strain yTL158, which expresses ORC with a halo-tagged orc3, was created by linearizing plasmid pRS303-CBP-TEV-halo-orc3-Gal1-10 orc4 with NheI and transforming it into yeast strain yTL151, which contains inducible expression plasmids for orc1, orc2, orc5, and orc6.

Labeling reactions: Proteins with HaloTag were labeled with JF646-HaloTag ligand (Promega # GA1120) by incubating the proteins with a tenfold excess of dye on ice for 0.5-1 h in the presence of 1 mM ATP. Free dye was removed by gel filtration (Superose 6 increase 10/300), and the labeling efficiency was determined to be 75% and 80% for JF646-ORC and JF646-MCM, respectively, after estimating protein and fluorophore concentrations relative to known standards. Accordingly, we cannot exclude the possibility that ∼25% and 20% of the observed single ORC or single MCM populations may have been partially labeled double ORC and double MCM hexamers.

Proteins with a single cysteine were labeled with Alexa Fluor 488 C5 Maleimide (Invitrogen # A10254) by incubating the proteins with a tenfold excess of dye on ice for 2 h at pH 7 in the absence of DTT. Free dye was removed by gel filtration (Superose 6 increase 10/300), for dCas9, or by dialysis with 2 M NaCl, 20 mM Tris-HCl pH 8, and 1 mM DTT followed by spin column chromatography (Zeba 7k MWCO) for histone octamers. Labeling efficiency was determined to be 81% for AF488-H2A, after estimating protein and fluorophore concentrations relative to known standards.

#### DNA substrates for single-molecule imaging

To generate a biotinylated 10.4 kbp DNA molecule containing ARS1 or mutated origin flanked by nucleosome positioning sequences, we ligated three different DNA fragments prepared by PCR. The left biotinylated arm with 6.6 kbp was amplified from plasmid pDRM1 (a kind gift from Daniel Ramírez-Montero) by PCR using primers HS_BN47 and HS_BN48 (ELLA Biotech) with Platinum SuperFi II DNA Polymerase (Thermo Scientific #12361010). The right biotinylated arm with 3.3 kbp was amplified as above from plasmid pDRM1 using primers HS_BN45 and HS_BN46 (ELLA Biotech). Biotinylated PCR products were purified by standard phenol-chloroform extraction, precipitated with ethanol and digested overnight with BsaI (NEB # R3733S). Digested biotinylated arms were purified by spin column chromatography using MicroSpin S-400 HR (Amersham # 27514001).

To generate a DNA fragment compatible for ligation containing ARS1 or mutated origin flanked by nucleosome positioning sequences, we first prepared template plasmids containing the origins. Plasmid template TL20-042 with ARS1 origin site flanked by 601 and a 603 nucleosome positioning sites^34^, was prepared by cloning into MluI-digested and Antarctic - dephosphorylated plasmid pSupercos1-lambda1,2^35^ of PCR fragment amplified from gBlock gene fragment (IDT) pTL013^16^ using primers TL-817 and TL-818, digested with AscI. Plasmid template TL22-072 with mutated ORC binding-site, was prepared by one-step cloning using NEBuilder HiFi reaction (NEB # E5520S) into plasmid pIA146^36^ linearized with HindIII (NEB # R3104S) of the fragment containing the mutated origin amplified from gBlock gene fragment (IDT)pGC218^23^ with primers TL-961 and TL-964, and the fragments containing the nucleosome positioning sites 601 and 603 independently amplified from gBlock gene fragment (IDT)pTL013^16^ with primers TL-958 – TL-959 and TL-962 – TL-963, respectively.

PCR fragments containing the origins and compatible BsaI ends were prepared from TL20-042 and TL22-072 by PCR using primers HS_BN23NPb and HS_BN26NPb (ELLA Biotech). PCR products were purified by standard phenol-chloroform extraction, precipitated with ethanol, digested overnight with BsaI (NEB # R3733S) and gel-purified.

Nucleosome assembly was carried out using salt gradient dialysis^37^. Fluorescently labeled histone octamers were mixed with DNA in High Salt Buffer (10 mM Tris pH 7.5, 2 M NaCl, 1 mM EDTA, 1 mM DTT). Samples were dialyzed for 18 h against 400 ml of High Salt Buffer and gradually supplemented with 2 L of Low Salt Buffer (10 mM Tris pH 7.6, 250 mM NaCl, 1 mM EDTA, 1 mM DTT). A final dialysis step for 1 h was performed into Zero Salt Buffer (20 mM Tris pH 7.5, 1 mM EDTA, 1 mM DTT). Fluorescently labeled histone octamer concentrations were optimized by small-scale titration and nucleosomes checked by 5% native PAGE. To test that the origin sequence is free of nucleosomes and accessible to ORC and MCM, the chromatinized construct were digested with Pst I (NEB#R0140S) and fragment size checked by 5% native PAGE. PCRs fragments with BsaI compatible ends were ligated to the chromatinized origins with T4 ligase overnight at 16 °C. The concentration of nucleosomes is maintained above 20 μg/ml to avoid dissociation due to dilution^38^ using commercial nucleosomes (Epicypher #160009). Final constructs were dialyzed overnight against 25 mM HEPES pH 7.6.

### Bulk Assays and Single-Molecule Experiments

#### MCM recruitment and loading reactions in bulk to test protein activity

Loading assays were carried out as follows: 50 nM ORC (or JF646-ORC), 50 nM Cdc6 and 100 nM Mcm2-7/Cdt1 (or JF646-Mcm2-7/Cdt1) were incubated with 300 ng DNA substrate (5.8 kbp circular bead-bound ARS1-containing pSK (+)-based plasmid^39^) coupled to magnetic beads for 30 min at 30 °C with mixing at 1250 rpm in 40 μl reaction buffer (25 mM HEPES-KOH pH 7.6, 10 mM MgOAc, 100 mM KOAc, 0.02% NP40, 5% glycerol, 1 mM DTT, and 5 mM ATP or ATPγS). Beads were then washed either with high salt wash buffer (45 mM HEPES-KOH pH 7.6, 5 mM MgOAc, 0.5 M NaCl, 0.02% NP-40, 10% glycerol, 1 mM EDTA, and 1 mM EGTA) followed by low salt wash buffer (45 mM HEPES-KOH pH 7.6, 5 mM MgOAc, 0.3 M KOAc, 0.02% NP-40, 10% glycerol, 1mM EDTA, and 1 mM EGTA), or only treated with low salt wash buffer. Finally, beads were resuspended in 10 μl elution buffer (45 mM HEPES-KOH pH 7.6, 5 mM MgOAc, 0.3 M KOAc, 10% glycerol, and 2 mM CaCl_2_), and DNA-bound proteins were released by MNase treatment (2 min 30° with 700 units of MNase NEB # M0247S) and analyzed by gel electrophoresis^40^.

#### Single-molecule instrumentation and visualization

A hybrid instrument combining optical tweezers and confocal microscopy was used to visualize the binding of DNA and protein at the single-molecule level (Q-Trap, LUMICKS) as described^7, 15^ with the following variations. The instrument makes use of a customer designed microfluidic flow cell with three inlets for injection of reaction buffers from the left and up to six inlets that are introduced orthogonally and can be used as protein reservoirs or buffer exchange locations in a temperature-controlled environment. Syringes and tubing connected to the flow cell were passivated, together with the flow cell itself, with 1 mg/ml BSA followed by 0.5% Pluronic F-127 (Sigma), each incubated for at least 30 min. Next, 20 pM of the biotinylated DNA, containing either a functional origin of replication or a mutated origin, chromatinized or not, was injected into one of the three laminar-flow-separated channels. Individual DNA molecules were trapped between two 1.76 μm diameter streptavidin-coated polystyrene beads (Spherotech) initially injected into a separate channel.

In all measurements, the stiffness of both optical traps was set to 0.3 pN/nm^41, 42^ The tethering of individual DNA molecules was verified by analysis of the force-extension curve obtained for each DNA molecule^43^ that was used for protein visualization. During fluorescence measurements, the DNA was held at a constant tension of 2 pN and the flow was turned off, unless otherwise specified. The AF488 and JF646 dyes were illuminated with two laser lines at 488 nm (2 µW) and 638 nm (7 µW), respectively, and the fluorescence from the dyes was detected on a single photon counting detector. Two-dimensional confocal scans were performed over an area of 90 x 18 pixels, which encompasses the DNA held at a force of 2 pN and the edges of both beads. The pixel size was set to 50×50 nm^2^, and the illumination time per pixel was set to 0.1 ms.

#### Protein concentrations and buffers in single-molecule experiments

Incubation and visualization of DNA-protein interactions in the flow cell were performed at 30 °C. ORC binding was conducted in reaction buffer (RB) containing 25 mM HEPES-KOH pH 7.6, 100 mM potassium glutamate, 10 mM magnesium acetate, 100 μg/mL BSA, 1 mM DTT, 0.01% NP-40-S, 10% glycerol, 5 mM ATP, with 10 nM Cdc6 and 5 nM JF646-ORC. To reduce the rate of photobleaching, we add 2 mM 1,3,5,7 cyclooctatetraene, 2 mM 4-nitrobenzylalchohol and 2 mM TROLOX.

Preparation of DNA-protein complexes in bulk for subsequent visualization in the flow cell was done as follows: 5 nM ORC was incubated with 1 nM 10.4 kbp biotinylated DNA (chromatinized or not) at 30 °C while mixing at 800 rpm in RB with 5 mM ATP. After 5 min, 10 nM Cdc6 was added to the reaction and incubated for a further 5 min. Then, 80 nM MCM/Cdt1 was added, bringing the total reaction volume to 50 μl. Following a 30 min incubation, samples were diluted 15 x in RB and injected into the microfluidic chip. ORC-Cdc6 DNA complexes were assembled in the same conditions while omitting Mcm2-7/Cdt1.

### Data analysis

#### Particle localization in 2D scans

We use the scikit-image (v0.16.2) implementation of a Laplacian of Gaussian (LoG) spot detector. The detection radius *r_LoG_* is set to 6.5 pixels (312 nm) for the red channel and 5 pixels (240 nm) for the blue channel; the LoG sigma parameter is given by *σ_LoG_* = *r_LoG_* / √2. We set the detection threshold to 0.2 ADU/pixel for the red channel and 0.3 ADU/pixel for the blue channel. For subpixel localization, detected spots are projected onto the x- and y-axes and then fitted with Gaussian profiles.

#### Particle tracking

The spots are tracked through subsequent frames using our own implementation of the Linear Assignment Problem method^44^. We used a maximum spot linking distance of 10 pixels (480 nm) for the red channel and 4 pixels (192 nm) for the blue channel using a maximum frame gap of 3 frames (1.8 seconds) without splitting. The LAP is solved with the scipy (v1.6.1) linear sum assignment optimizer^45^. Two spots are considered colocalized if they are on average less than 4 pixels (200 nm) apart over the first 5 frames; spot intensities are calculated by taking the total ADU count within the detection radius.

#### Location calibration

Using a reference dCas9 dataset, we map the location of two known DNA sequences onto pixel coordinates, relative to the left bead (whose coordinate is given in microns by the C-Trap metadata, as measured in the brightfield view). With these pixel coordinates we calculate the inverse transformation, from pixel coordinate to DNA sequence location in base pairs, giving us a brightfield-to-confocal offset value (4.5 pixels) and a pixel size value (48 nm per pixel).

#### Fluorophore intensity calibration

The reference dCas9 dataset is also used for fluorophore intensity calibration. Foci with one bleaching step are used to make a distribution of intensity bleaching steps and calculate the mean *μ_ΔI_* and standard deviation *σ_ΔI_* of that distribution. The minimum bleaching step size, which is needed for stoichiometry determination of further experiments, is set to *ΔI_min_ ≤ μ_ΔI_ - 2 σ_ΔI_* in order to capture at least 95% of all bleaching events. The measured values of *μ_ΔI_* and *σ_ΔI_* for each color are given in **Table 1**.

**Table 1.** Fluorophore properties, calibrated using dCas9 data.

| Fluorophore | Signal | $\mu_{\Delta I}$ (ADU) | $\sigma_{\Delta I}$(ADU) | $\Delta I_{min}$ (ADU) | $N_{foci}$ |
| --- | --- | --- | --- | --- | --- |
| AF488 | blue | 41.6 | 9.8 | 20 | 43 |
| JF646 | red | 176.7 | 22 | 75 | 14 |

#### Stoichiometry determination

To determine the number of fluorophores present within each detected spot, we perform photobleaching step counting using Change-Point Analysis (CPA). We use the ruptures (v1.1.6) ^46^ implementation with an L2 cost function to detect mean-shifts in the signal. The minimum segment length is set to 2 and the penalty is set to *ΔI* ^2^. If any steps larger than *ΔI_min_* are present after the fit, the smallest steps are combined together (“pruned”) until only steps larger than *ΔI_min_* are left.

#### Data filtering

The resulting data table of traces with number of fluorescent proteins per spot was filtered in order to reduce noise, outliers, and data that is not suitable for further motion analysis: Diffraction-limited spots containing more than 5 fluorescent proteins, likely aggregates, are filtered out.

Any traces starting or ending within 1 kbp from a bead are filtered out to prevent any proteins likely stuck to a bead from entering the dataset.

Any traces starting after frame 5 are also filtered away because we do not expect any fluorescent protein to land on the DNA during the scan.

Finally, only traces with a length of 5 frames or more are retained for diffusion analysis.

#### Spatial distribution analysis

In spatial distribution plots we show the average position of the spot over the first three frames. The bin size of the histogram is set to 670 bp to be close to (but slightly larger than) the diffraction limit. Together with the binned data we plot the kernel density estimation of the data with a bandwidth that is equivalent to half of the bin size in the histogram (335 bp) and is higher than the localization error in the imaging conditions (100 bp).

#### Motion analysis

We analyzed the mean squared displacement (MSD) of individual tracked foci as a function of the delay time between frames as previously described^15, 47^. We employ a Gaussian Mixture Model to fit the distribution of log(*D*) in the data set without a chromatinized origin, in order to differentiate between different kinetic populations. We assess the statistical preference for either a two-state or single-state model using the Bayesian Information Criterion, and the two-state model is favored. We identified these states as a static population (*D_slow_* = 0.0049 ± 0.0028 kbp^2^ s^-1^ (mean ± SEM)) and a diffusive population (*D_fast_* = 0.152 ± 0.023 kbp^2^ s^-1^). The means of these subpopulations are imposed in the fits to the other datasets.

We calculated the mean, variance, and standard error (*m, var, SEM*) of the *i^th^*diffusion coefficient distribution from the fitted lognormal parameters *µ_i_* and *σ_i_* according to:

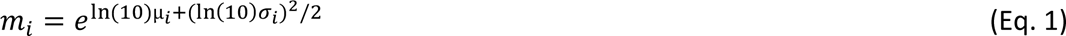

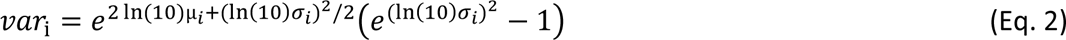

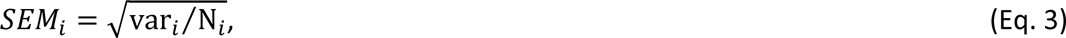

where the factors of log(*e*) and ln(10) account for our use of base 10, and *N_i_* is the number of values in the data set times the area of the *i^th^* fitted peak. It is worth noting that the dependence of *m_i_* on both *µ_i_* and *σ_i_* yields mean diffusion constants larger than might be inferred by simply computing 10^μ*i*^.

#### Probability model for H2A stoichiometry distributions

We formulate a probability model for H2A stoichiometry based on three probabilistic quantities, the site occupation probability *P_occ_*, the H2A binding probability *P_H2A_*, and the labeling efficiency *P_lab_*.

The probability of having 0, 1, or 2 occupied sites (*N_occ_*) on the DNA depends on the site occupation probability (*P_occ_*) following the binomial distribution:

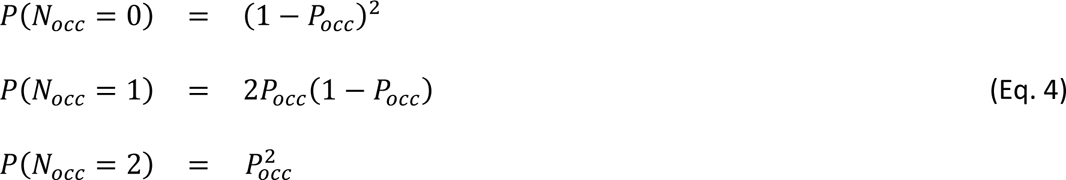

An H2A-H2B dimer binds to the tetrasome with a probability of *P*_*H*2*A*_. Again following the binomial distribution, the probabilities of a tetrasome, hexasome and full nucleosome forming are respectively:

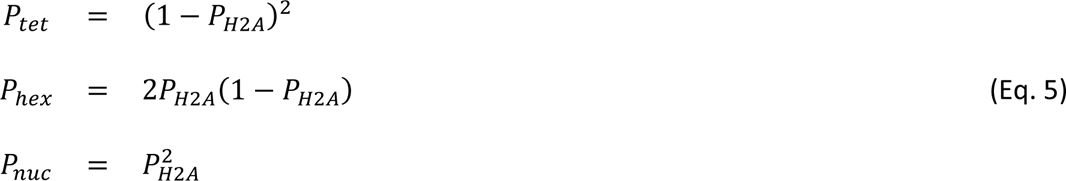

Combining the two parts listed above yields the probabilities for encountering the possible numbers of H2A (*N*_*H*2*A*_) on the DNA:

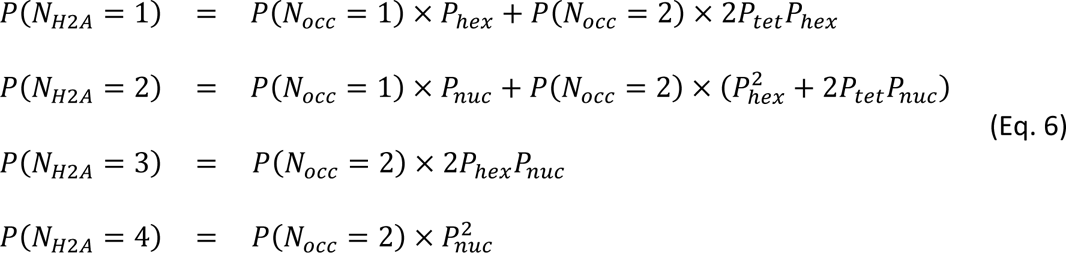

Finally, we need to take labeling efficiency *P_lab_* into account. This gives us the probabilities for finding *N*_*vis*_ visible fluorophores in a diffraction-limited spot on the DNA:

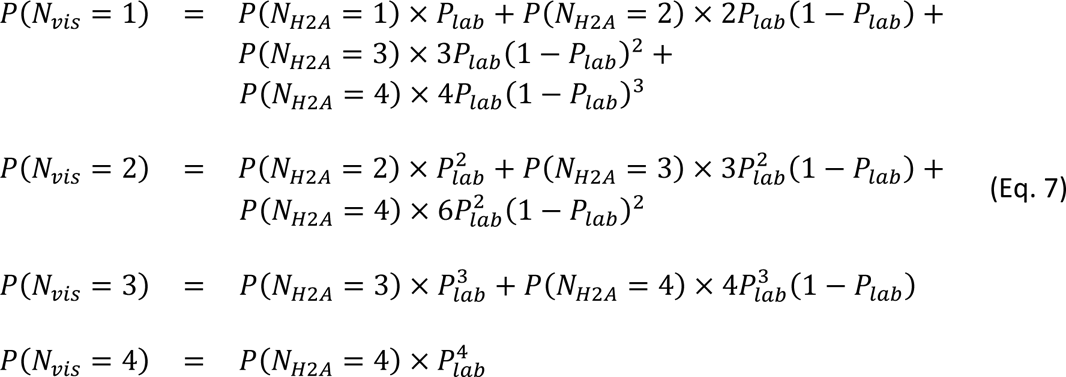

#### Force analysis

To quantify unwrapping of DNA from nucleosomes, we performed a transformation of force-distance curves to contour length space. The persistence length *L_p_* and stretch modulus *S* of dsDNA in the measurement buffer were determined by pulling on bare dsDNA molecules to the overstretching regime at a constant pulling speed of 100 nm/s and fitting the force-distance curves using the extensible worm-like chain model^48^:

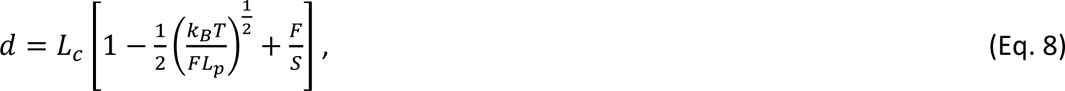

where *d* is the distance between the two ends of the dsDNA, *L_c_* is the DNA contour length, *k_B_* is Boltzmann constant, and *F* is the pulling force. The fitting results are *L_p_* = 44.2 nm and *S* = 1421 pN.

The force-distance curves obtained by pulling on the chromatin sample were transformed to contour length space by calculating the DNA contour length *L_c_* using Eq. 9, an inversion of Eq. 8, and plotting *L_c_* against pulling force applied to the chromatin:

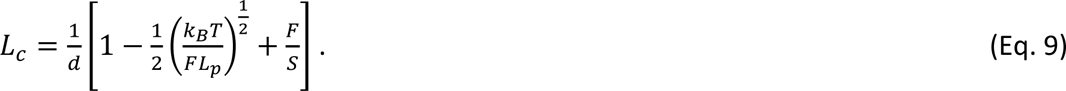

This results in graphs with segments of constant *L_c_*. CPA (again using the ruptures library with an L2 cost function) was applied to the *L_c_*-*F* plots to identify the plateaus and extract contour length increments between the plateaus, which resulted from unwrapping of nucleosomes, as well as the nucleosome unwrapping forces. This process follows the same steps as the fluorophore bleaching step fitting analysis, but here we set the minimum step size to 10 nm, after calibration with dsDNA.

#### Experiment automation

To automate some of the C-trap measurements, we used the Lumicks Harbor experiment automation scripts (https://harbor.lumicks.com/scripts) as a starting point; most notably, Joep Vanlier’s automation script for catching beads, fishing for DNA, and making force-distance curves. We added the functionality to automatically acquire confocal images after successfully trapping DNA.

#### Error determination

To compute the statistical error in population, stoichiometry and diffusion plots, we use the Wilson confidence interval. This is an improvement over the normal approximation for the error of sample proportion, especially for a small number of trials^49^. We use the statsmodels (Python) implementation for its calculation^50^.

#### Figure schematics

Figure schematics were generated using BioRender.com (Standard Academic License).

#### Data availability

The data that support the findings of this study are available from the authors upon reasonable request.

#### Code availability

The analysis of the acquired data was performed using custom-written scripts in Python 3.8, which are available upon request.

## ACKNOWLEDGMENTS

We thank Anne Early and Lucy Drury for providing yeast strains, Nerea Murugarren for assistance in loading bulk biochemical assays, Fiona Horne and Emilie van Vet for preliminary chromatin assembly experiments, and Alessandro Costa for useful discussions. ZL acknowledges the support given by EMBO Postdoctoral Fellowship (ALTF 484-2022). ND acknowledges funding from the Netherlands Organization for Scientific Research (NWO) through Top grant 714.017.002), ‘BaSyC—Building a Synthetic Cell’ Gravitation grant (024.003.019) of the Netherlands Ministry of Education, Culture and Science (OCW), and from the European Research Council through an Advanced Grant (REPLICHROMA; grant number 789267.

## AUTHOR CONTRIBUTIONS

HS, JD, and ND conceived the study, and ND supervised the study. HS designed the experiments with input from ZL, EvV, JD, and ND. JD provided cell strains and advised on protein purification and biochemical conditions. TvL engineered DNA substrates, purified the proteins, and performed bulk biochemical assays. HS labeled the proteins and assembled chromatin substrates. HS performed all single-molecule experiments. ZL and EvV designed and wrote data analysis routines. HS and EvV also performed data analysis, with input from ZL and ND. All authors were involved in discussion of the data. HS and ND wrote the manuscript with input from the other authors.

