## Supplementary_figures for "A chromatinized origin reduces the mobility of ORC and MCM through interactions and spatial constraint" (from another version of this preprint: version 2, 2023-06-26)

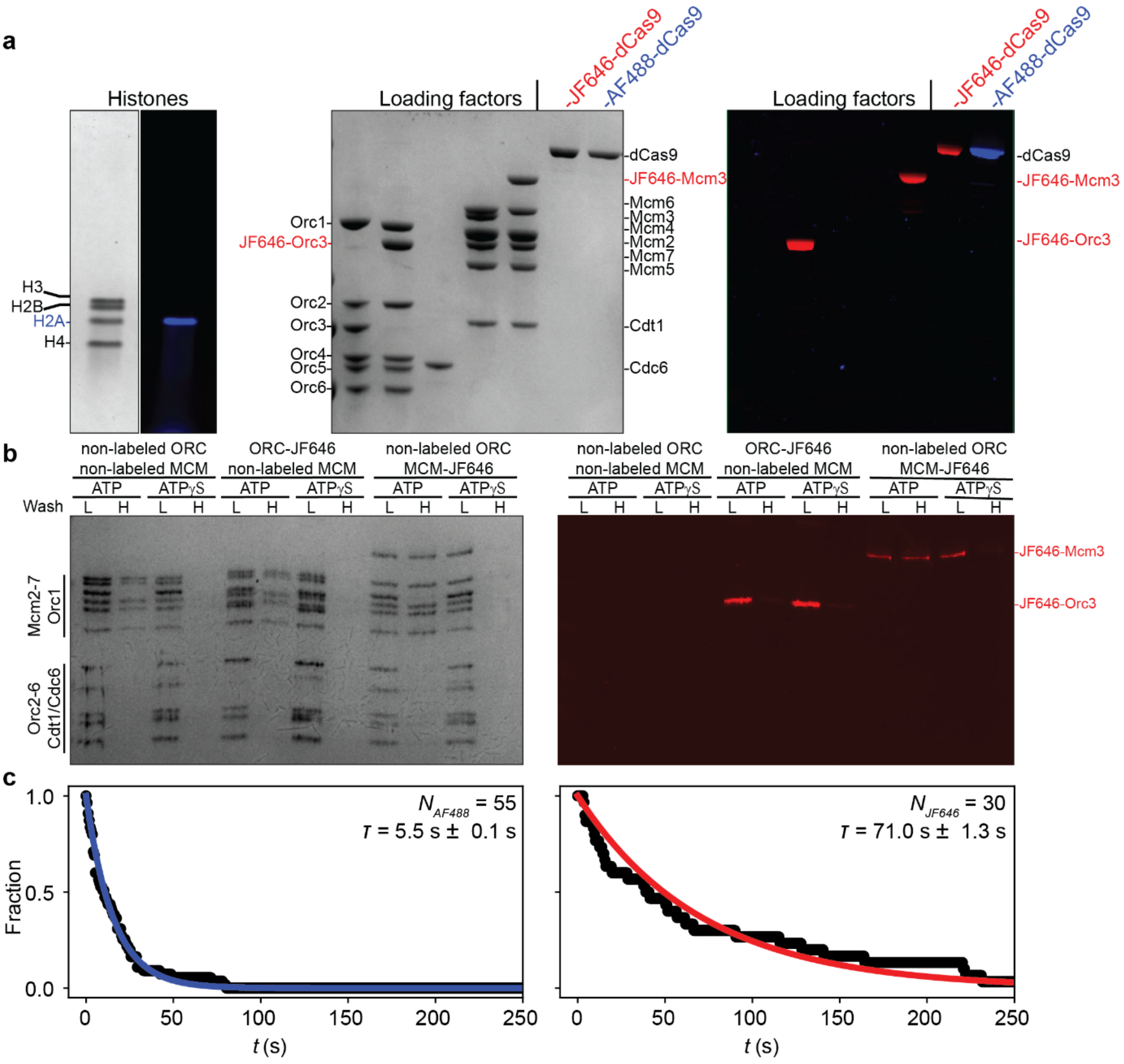


**Supplementary Figure 1.1. Protein purification and ensemble characterization (a)** Scans from the SDS-PAGE gels after Coomassie staining (in b/w) and after fluorescence imaging (in color) of the yeast histone octamer complex (containing labeled AF488-H2A), loading factors (ORC complex, labeled ORC complex, including a HaloTag on the Orc3 subunit labeled with a single JF646 dye, Cdc6, Mcm2-7/Cdt1 complex, labeled Mcm2-7/Cdt1 complex, including a HaloTag on the Mcm3 subunit labeled with a single JF646 dye) and fluorescence standards dCas9 labeled with Halo-JF646 and Halo-AF488. **(b)** Loading assay performed on bead-tethered DNA in bulk using non-labeled proteins, labeled ORC complex with non-labeled Mcm2-7/Cdt1 and non-labeled ORC complex with labeled Mcm2-7/Cdt1. Following a reaction performed in the presence of either ATP or ATPγS, either a low (L) or a high-salt (H) wash is performed. DNA is removed from the beads by MNase treatment, and the presence of DNA-bound proteins is examined using SDS-PAGE followed by fluorescence scanning (color image) and silver staining (b/w). Mcm2-7 proteins are retained on the DNA following a high-salt wash only if the loading reaction was performed in the presence of ATP. This confirms recruitment of Mcm2-7 in the presence of ATPγS, but loading of Mcm2-7 only in the presence of ATP. **(c)** Lifetime analysis of the fluorophores used in this study obtained with an acquisition frequency of one frame every 0.6 s. (left panel) Survival probability of single AF488 linked to dCas9 ($N_{AF488}$). (right panel) Survival probability of single JF646 linked to dCas9 ($N_{JF646}$). The black dots represent the data, and the solid curves are exponential decay fits to the data. The error shown is the standard deviation, computed from the covariance of the optimal parameters found by the least squares fit.

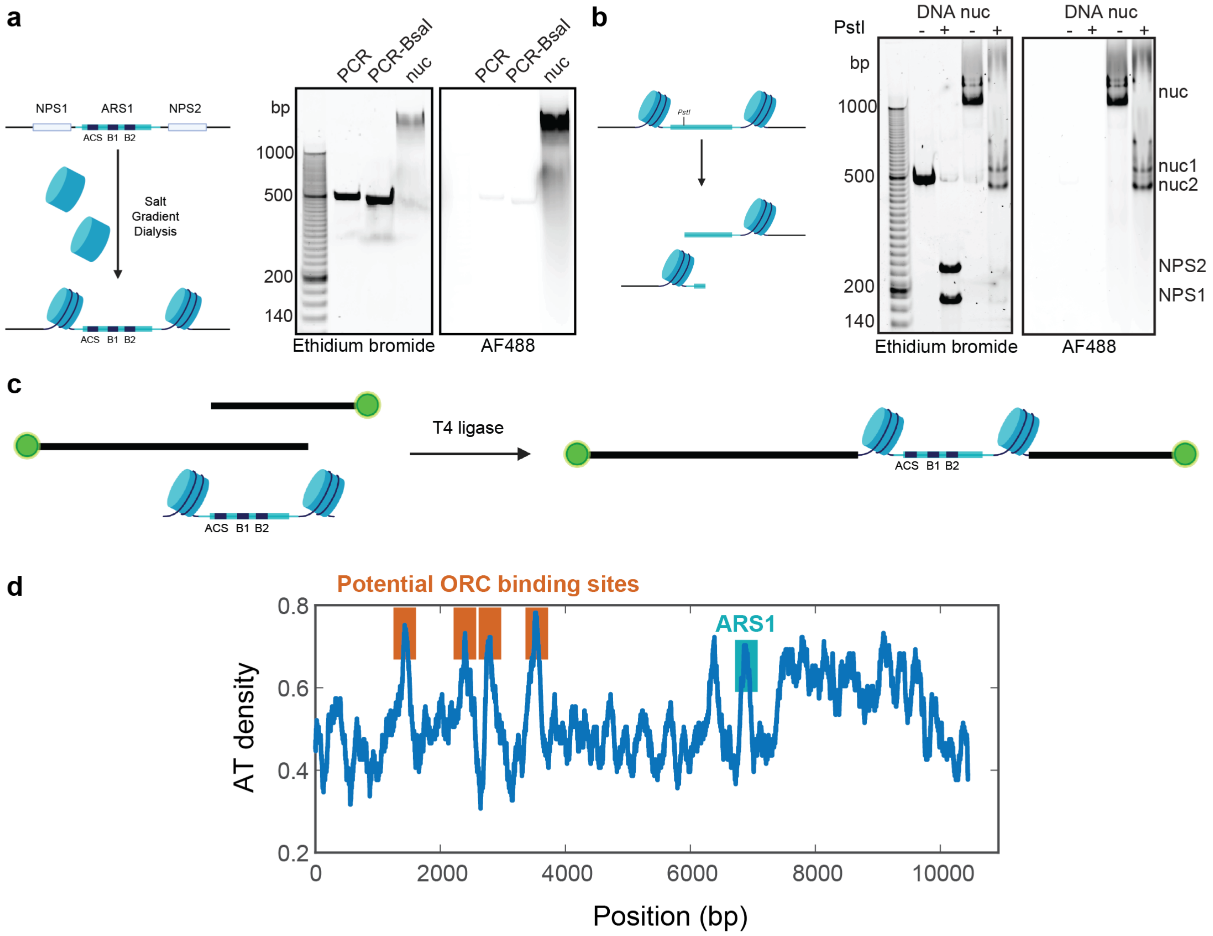


**Supplementary Figure 1.2. Establishment of a chromatinized origin on a 10.4 kbp DNA. (a)** (left panel) Schematic of a PCR fragment that contains the ARS1 origin of replication flanked by two NPSs used for nucleosome assembly by salt gradient dialysis. (right panel) Scans from a native PAGE gel after ethidium bromide staining (left) and after fluorescence imaging (right) of the PCR fragment, PCR fragment with cohesive ends, and PCR fragment with cohesive ends with assembled nucleosomes prior to ligation. **(b)** (left panel) Schematic of the restriction enzyme accessibility assay performed to determine the degree of NPS occupancy. (right panel) Scans from a native PAGE gel after ethidium bromide staining (left) and after fluorescence imaging (right) of the PCR with cohesive ends (DNA) without (-) or with (+) PstI, and nucleosomes assembled on the DNA (nuc) without (-) or with (+) PstI. **(c)** Schematic of the ligation protocol using T4 ligase to generate the final 10.4 kbp DNA from the assembled nucleosomes and the biotinylated ‘arms’. **(d)** Nucleotide density analysis along the 10.4 kbp DNA construct in sliding windows of 100 bp. ORC near-cognate binding sites and the ARS1 site position are indicated in the plot.

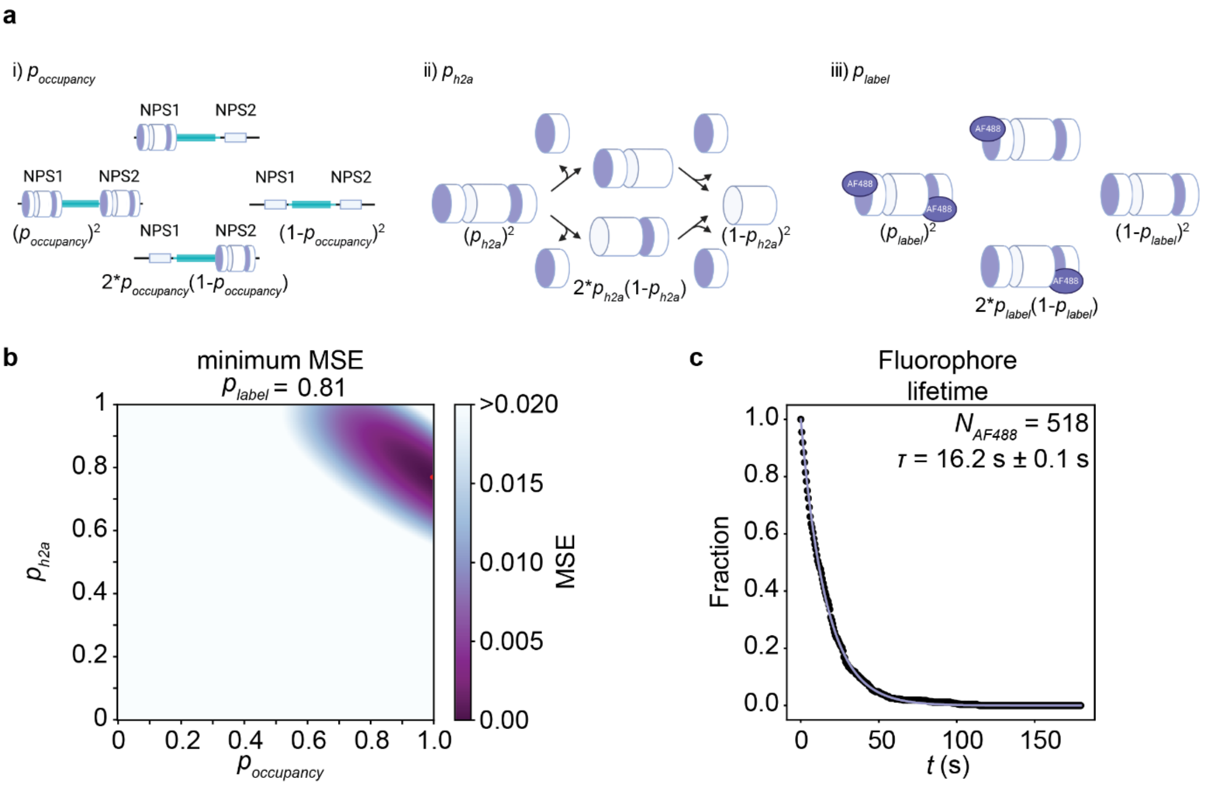


**Supplementary Figure 1.3. Model for extracting biophysical parameters from the H2A stoichiometry distributions. (a)** Three factors affect the visibility of fluorescently labeled H2A in our experiments: (i) Chromatinized molecules can have two, one or zero nucleosomes ($p_{occupancy}$, free parameter); (ii) H2A-H2B dimers can dissociate from the full nucleosomes, resulting in either hexasomes with one fluorophore or unlabeled H3-H4 tetrasomes ($p_{h2a}$, free parameter). (iii) The labeling efficiency of H2A (experimentally determined and input into the model as $p_{label}$) can result in unlabeled nucleosomes, nucleosomes with one label or nucleosomes with two labels; **(b)** Landscape of the mean squared error (MSE) between fit and experiment, calculated for different values of $p_{h2a}$ and $p_{occupancy}$ at a fixed value of $p_{label}$ = 0.81. The red dot indicates the optimal solution with the lowest error ($p_{h2a}$ = 0.77, $p_{occupancy}$ = 1.00). **(c)** Lifetime analysis of H2A. Survival probability of all ($N_{AF488}$ = 518) single AF488 associated to H2A on DNA containing an origin as deduced from the diffraction-limited blue spots collected from 248 distinct DNA molecules with an acquisition frequency of one frame every

0.6 s. The error shown is the standard deviation, computed from the covariance of the optimal parameters found by the least squares fit.

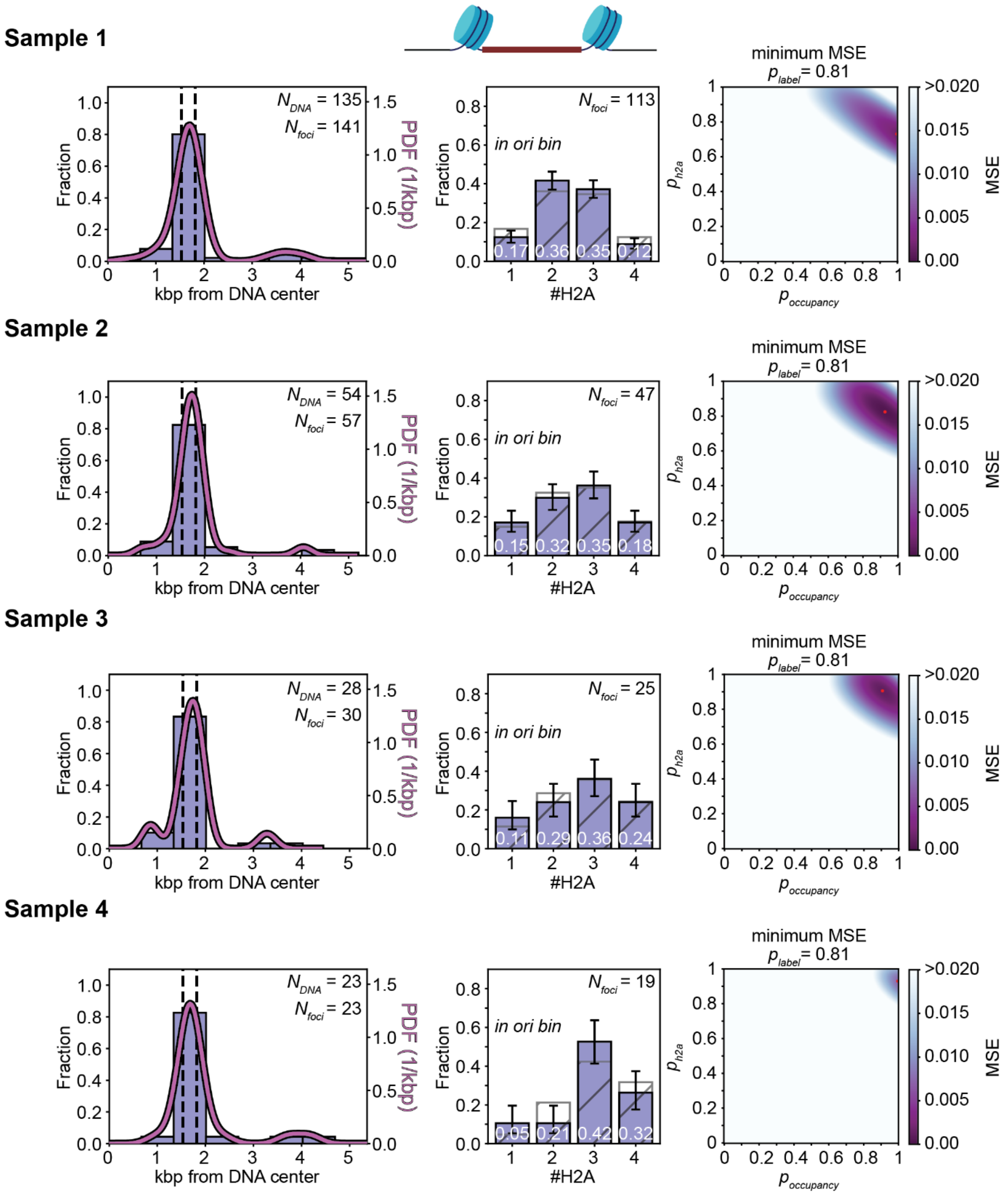


**Supplementary Figure 1.4. Variability between sample preparation of a labeled chromatinized origin within 10.4 kbp DNA.** (left panels) Spatial distribution of fluorescently labeled H2A on DNA as described in **Figure 1b**, deduced from the blue diffraction-limited spots ($N_{foci}$) collected from a number of DNA molecules indicated by $N_{DNA}$. Dashed lines indicate the location of the NPSs, and the solid curve indicates the kernel density estimation of the data. (center panels) Stoichiometry distribution of fluorescently labeled H2A histones in the bin containing the chromatinized origin. Error bars are one-sigma Wilson confidence intervals. White numbers inside the bars designate the fitted values based on the model described in the Methods and in **Supplementary Figure 1.3**. (right panels) Landscapes of the mean squared error (MSE) between fit and experiment, calculated for different values of $p_{h2a}$ and $p_{occupancy}$ at a fixed value of $p_{label}$ = 0.81. The red dots indicate the optimal solutions with the lowest error.

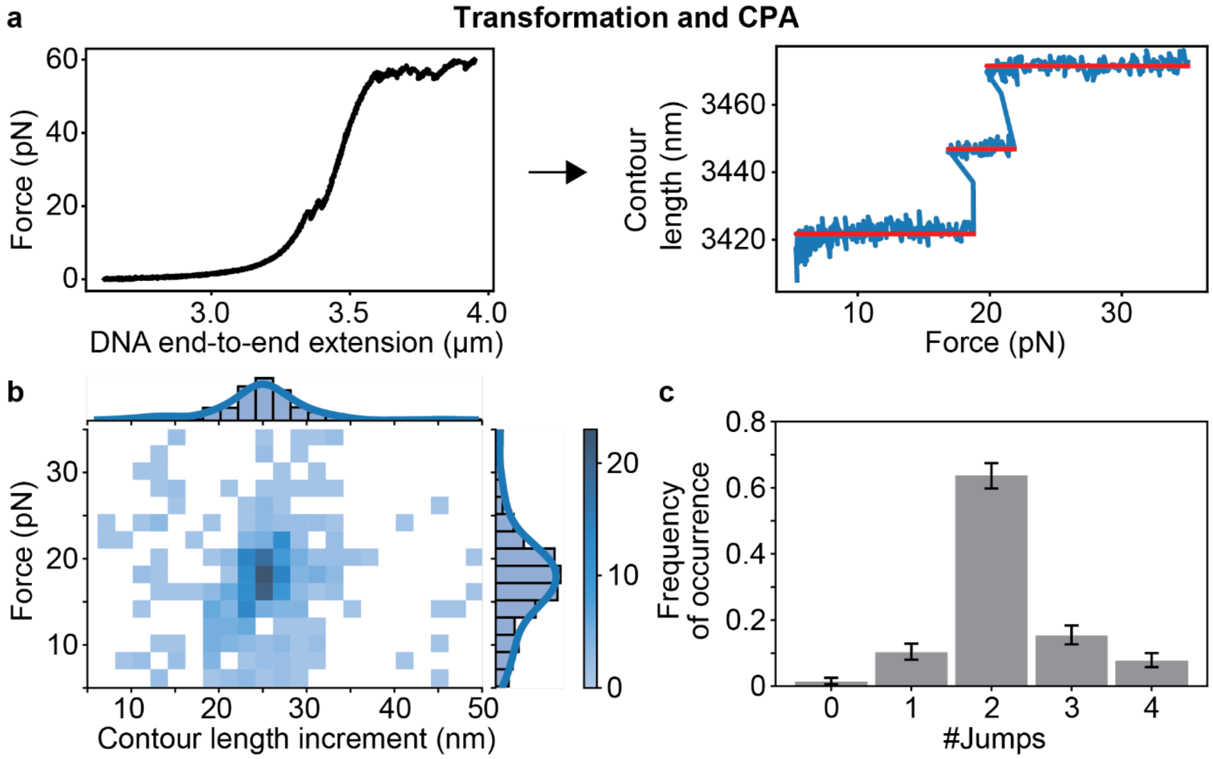


**Supplementary Figure 1.5. Force spectroscopy of 10.4 kbp DNA containing a chromatinized origin. (a)** (left panel) Representative force-extension curve for a typical chromatinized DNA. Unwrapping of the DNA from the nucleosomes results in observable increases in the end-to-end extension of the DNA and concomitant decreases in the force[23]. The extent of DNA unwrapped from the nucleosomes is assessed by transforming the data into a contour length-force plot and performing change point analysis (CPA) to detect the plateaus. **(b)** Heatmap of the distribution of force (<$F_{unwrap}$> = 17.6 pN ± 5.8 pN) and contour length increments (<$\Delta L_c$> = 24.8 nm ± 5.9 nm) found by CPA in 157 molecules. **(c)** Distribution of the number of contour length increments per chromatinized DNA molecule.

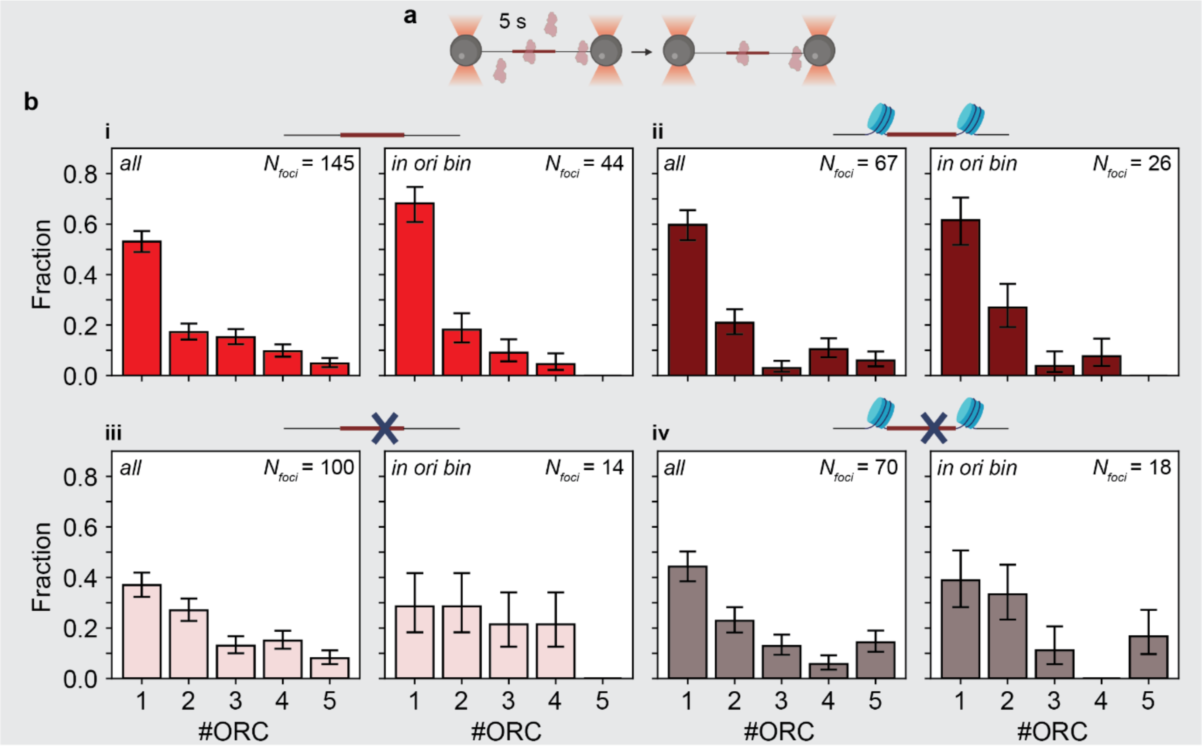


**Supplementary Figure 2. Stoichiometry of ORC on 10.4 kbp DNA containing a chromatinized origin. (a)** Incubation method pertaining to data in panel b. Tethered DNA molecules (chromatinized or not, as indicated) are introduced into Channel 4 for incubation with ORC and Cdc6 for 5 s in the presence of ATP, and then moved into the buffer channel (Ch3) for confocal scanning. **(b)** Stoichiometry of ORC foci ($N_{foci}$) from data shown in **Figure 2.** (left panels) Stoichiometry distributions of ORC over the entire DNA molecule and (right panels) stoichiometry distribution of ORC in the bin containing (i) the non-chromatinized ARS1 origin; (ii) the chromatinized ARS1 origin; (iii) the non-chromatinized mutated origin; and (iv) the chromatinized mutated origin, as also indicated by the schematics.

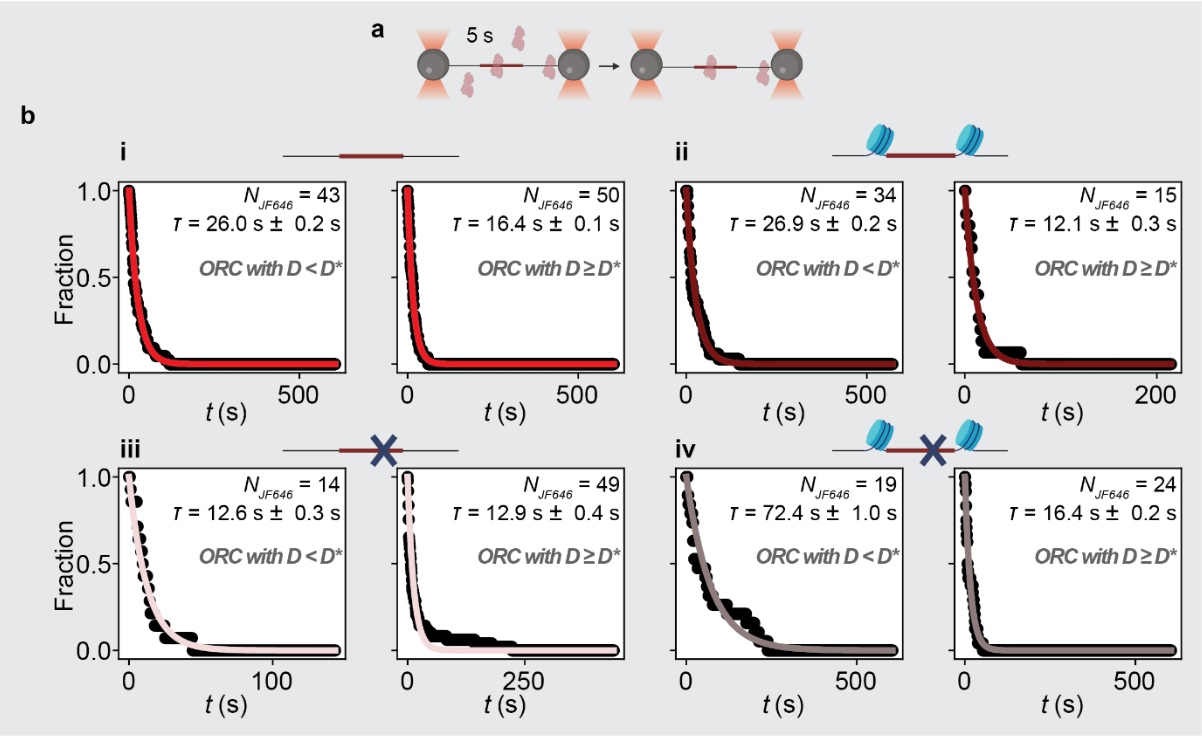


**Supplementary Figure 3. Lifetime of ORC on 10.4 kbp DNA containing a chromatinized origin. (a)** Incubation method pertaining to data in panel b. Tethered DNA molecules (chromatinized or not, as indicated) are introduced into Channel 4 for incubation with ORC and Cdc6 for 5 s in the presence of ATP, and then moved into the buffer channel (Ch3) for confocal scanning with an acquisition frequency of one frame every 0.6 s. **(b)** (i) (left panel) Survival probability of single JF646 ($N_{JF646}$ = 43) associated to slowly moving or static ORC on DNA containing an origin as deduced from the diffraction-limited red spots with $D < D^*$. (right panel) Survival probability of single JF646 ($N_{JF646}$ = 50) associated to diffusive ORC on DNA containing an origin as deduced from the diffraction-limited red spots with $D \geq D^*$. (ii) Survival probability of single JF646 ($N_{JF646}$ = 34) associated to slowly moving or static ORC on DNA containing an origin as deduced from the diffraction-limited red spots with $D < D^*$. (right panel) Survival probability of single JF646 ($N_{JF646}$ = 15) associated to fast diffusive ORC on DNA containing an origin as deduced from the diffraction-limited red spots with $D \geq D^*$. (iii) (lower left panel) Survival probability of single JF646 ($N_{JF646}$ = 14) associated to slowly moving or static ORC on DNA containing an origin as deduced from the diffraction-limited red spots $D < D^*$. (right panel) Survival probability of single JF646 ($N_{JF646}$ = 49) associated to fast diffusive ORC on DNA containing an origin as deduced from the diffraction-limited red spots with $D \geq D^*$. (iv) (lower left panel) Survival probability of single JF646 ($N_{JF646}$ = 19) associated to slowly moving or static ORC on DNA containing an origin as deduced from the diffraction-limited red spots with $D < D^*$. (right panel) Survival probability of single JF646 ($N_{JF646}$ = 24) associated to fast diffusive ORC on DNA containing an origin as deduced from the diffraction-limited red spots with $D \geq D^*$. The error shown is the standard deviation, computed from the covariance of the optimal parameters found by the least squares fit.

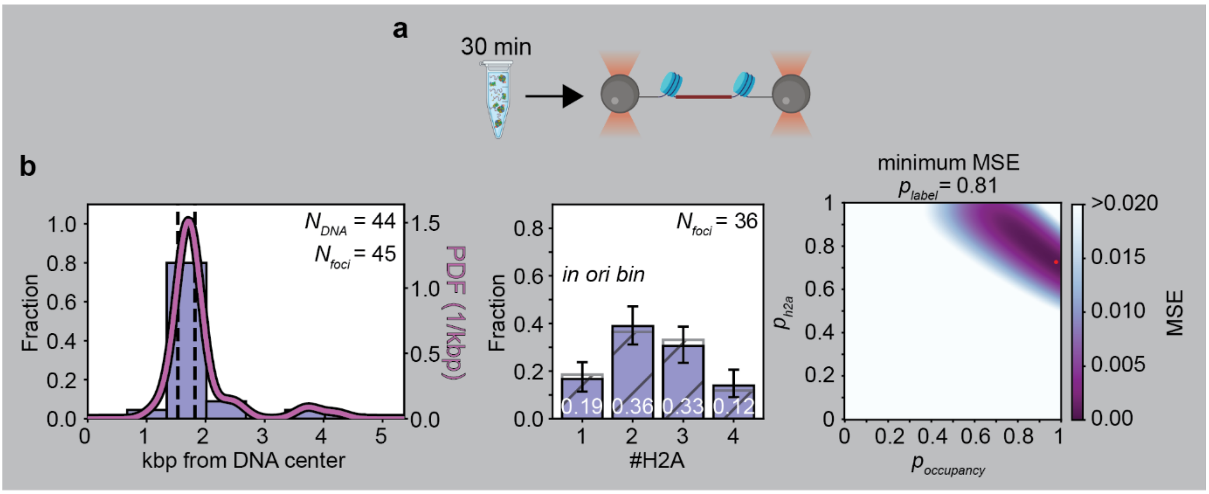


**Supplementary Figure 4.1. Stability of the labeled chromatinized origin on 10.4 kbp DNA in MCM loading buffer. (a)** DNA molecules containing a chromatinized ARS1 origin are incubated in bulk for 30 min in the presence of ATP. Subsequently, the DNA-protein construct is flushed into the single-molecule flow cell, tethered and moved into the buffer channel for confocal scanning. **(b)** (left panel) Spatial distribution of fluorescently labeled H2A on DNA as described in **Figure 1b**, deduced from the blue diffraction-limited spots ($N_{foci}$) collected from a number of DNA molecules indicated by $N_{DNA}$. Dashed lines indicate the location of the NPSs, and the solid curve indicates the kernel density estimation of the data. (center panel) Stoichiometry distribution of fluorescently labeled H2A histones in the bin containing the chromatinized origin. Error bars are one-sigma Wilson confidence intervals. White numbers inside the bars designate the fitted values based on the model described in the Methods and in **Supplementary Figure 1.3**. (right panel) Landscape of the mean squared error (MSE) between fit and experiment, calculated for different values of $p_{h2a}$ and $p_{occupancy}$ at a fixed value of $p_{label}$ = 0.81. The red dot indicates the optimal solution with the lowest error. Data derives from one chromatinized sample.

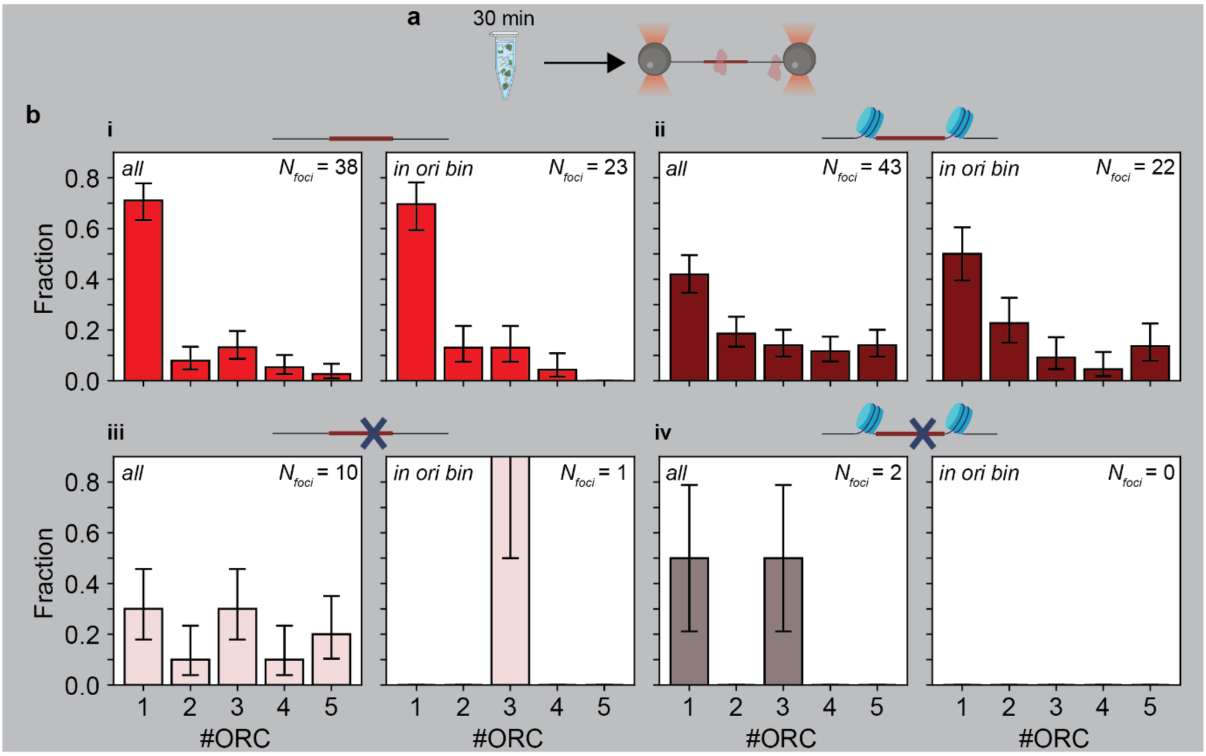


**Supplementary Figure 4.2. Stoichiometry of ORC on 10.4 kbp DNA containing a chromatinized origin. (a)** Incubation method pertaining to data in panel b. ORC and Cdc6 are incubated with DNA molecules (chromatinized or not, as indicated) in bulk for 30 min in the presence of ATP. Subsequently, the DNA-protein construct is flushed into the single-molecule flow cell, tethered, and moved into the buffer channel for confocal scanning. **(b)** Stoichiometry of ORC foci ($N_{foci}$) from data shown in **Figure 4**. (left panels) Stoichiometry distributions of ORC over the entire DNA molecule and (right panels) stoichiometry distribution of ORC in the bin containing (i) the non-chromatinized ARS1 origin; (ii) the chromatinized ARS1 origin; (iii) the non-chromatinized mutated origin; and (iv) the chromatinized mutated origin, as also indicated by the schematics.

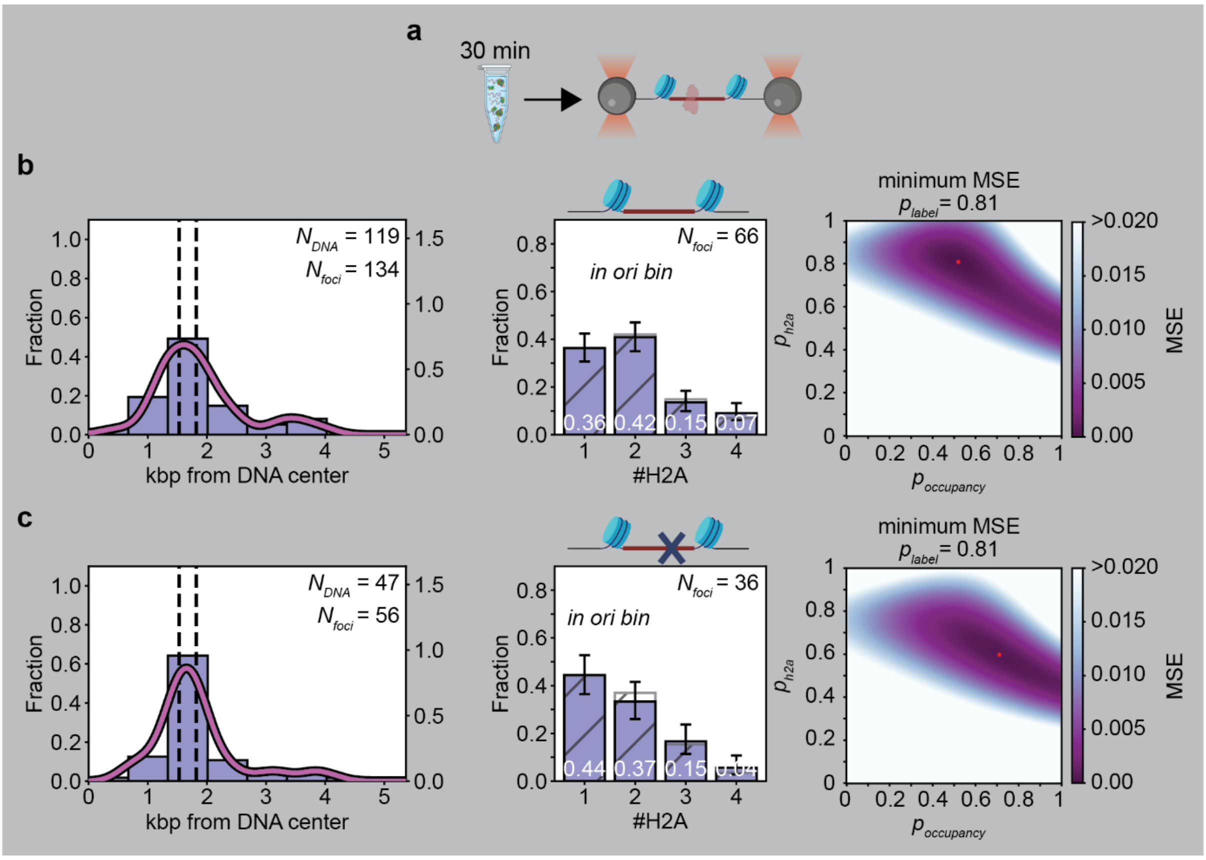


**Supplementary Figure 4.3. Stability of the labeled chromatinized origin on 10.4 kbp DNA in the presence of ORC and Cdc6. (a)** Incubation method pertaining to data in panels (b-c). ORC and Cdc6 are incubated with chromatinized DNA molecules in bulk for 30 min in the presence of ATP. Subsequently, the DNA-protein construct is flushed into the single-molecule flow cell, tethered, and moved into the buffer channel for confocal scanning. **(b)** (left panel) Spatial distribution of fluorescently labeled H2A on DNA containing a chromatinized ARS1 origin as described in **Figure 1b**, deduced from the blue diffraction-limited spots ($N_{foci}$) collected from a number of DNA molecules indicated by $N_{DNA}$. Dashed lines indicate the location of the NPSs, and the solid curve indicates the kernel density estimation of the data. (center panel) Stoichiometry distribution of fluorescently labeled H2A histones in the bin containing the chromatinized origin. Error bars are one-sigma Wilson confidence intervals. White numbers inside the bars designate the fitted values based on the model described in the Methods and in **Supplementary Figure 1.3**. (right panel) Landscape of the mean squared error (MSE) between fit and experiment, calculated for different values of $p_{h2a}$ and $p_{occupancy}$ at a fixed value of $p_{label}$ = 0.81. The red dot indicates the optimal solution with the lowest error. Data derives from one chromatinized sample. **(c)** (left panel) Spatial distribution of fluorescently labeled H2A on DNA containing a chromatinized mutated origin as described in panel (b). (center panel) Stoichiometry distribution of fluorescently labeled H2A histones in the bin containing the chromatinized mutated origin. (right panel) Landscape of the mean squared error (MSE) between fit and experiment, calculated as in panel (b). Data derives from one chromatinized sample.

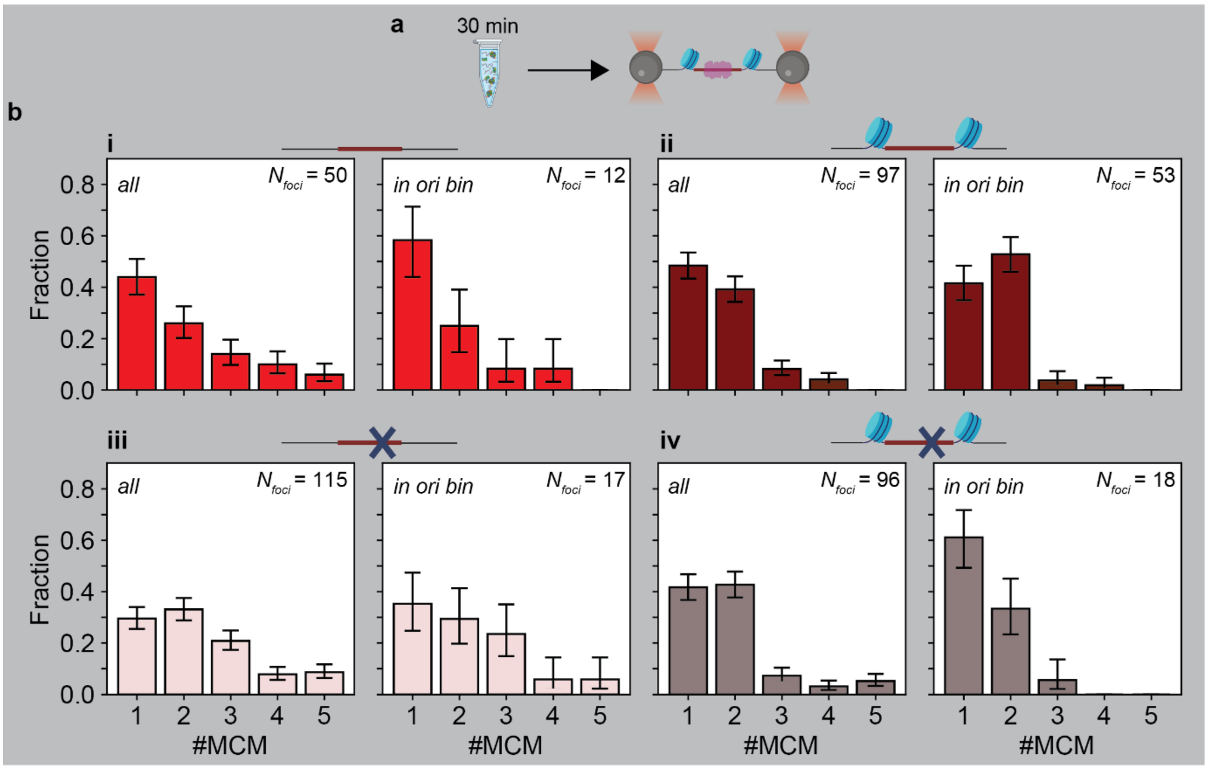


**Supplementary Figure 5.1. Stoichiometry of MCM on 10.4 kbp DNA containing a chromatinized origin. (a)** ORC, Cdc6, and Mcm2-7/Cdt-1 are incubated with 10.4 kb DNA molecules (chromatinized or not, as indicated) in bulk for 30 min in the presence of ATP. Subsequently, the DNA-protein construct is flushed into the single-molecule flow cell, tethered and moved into the buffer channel for confocal scanning. **(b)** Stoichiometry of MCM foci ($N_{foci}$) from data shown in **Figure 4.** (left panels) Stoichiometry distributions of MCM over the entire DNA molecule and (right panels) stoichiometry distribution of MCM in the bin containing (i) the non-chromatinized ARS1 origin; (ii) the chromatinized ARS1 origin; (iii) the non-chromatinized mutated origin; and (iv) the chromatinized mutated origin, as also indicated by the schematics.

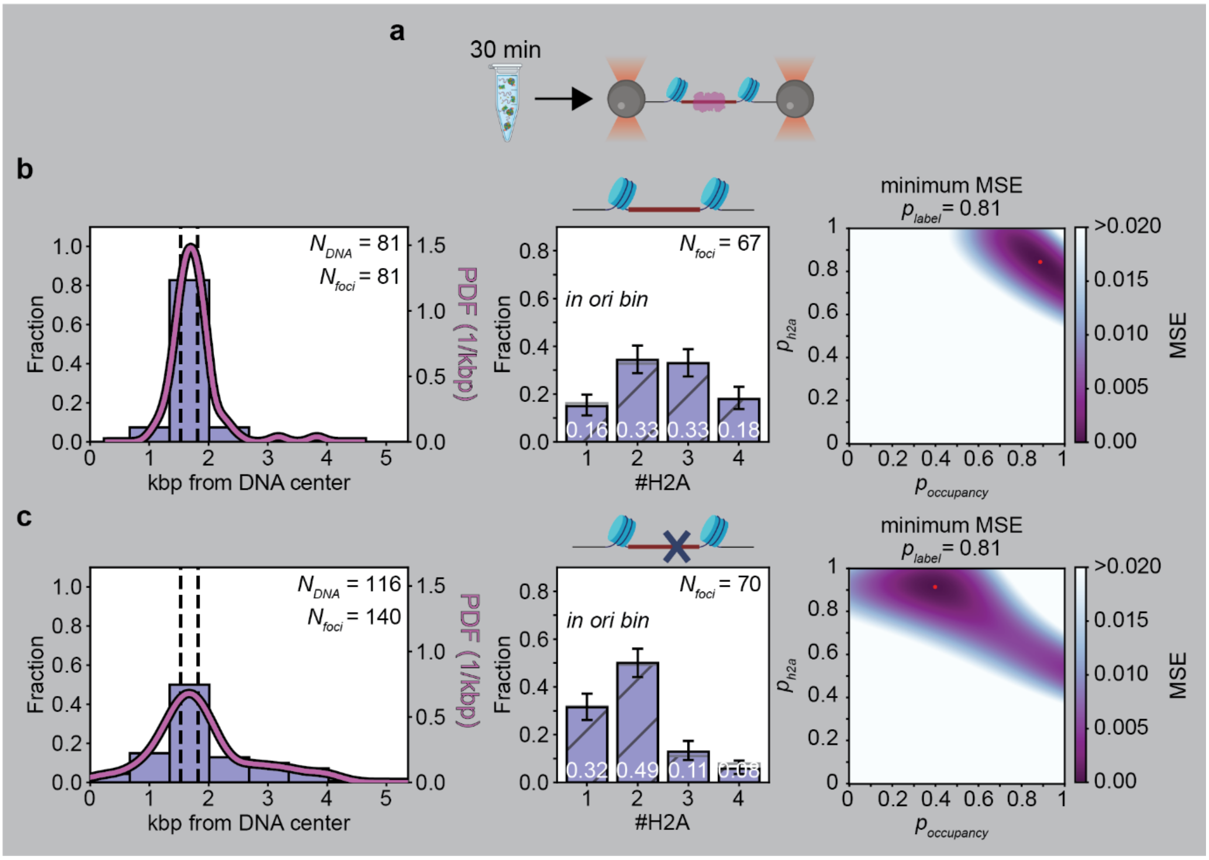


**Supplementary Figure 5.2. Stability of the labeled chromatinized origin on 10.4 kbp DNA in the presence of MCM, ORC and Cdc6. (a)** Incubation method pertaining to data in panels (b-c). ORC, Cdc6, and Mcm2-7/Cdt-1 are incubated with 10.4 kb DNA molecules (chromatinized or not, as indicated) in bulk for 30 min in the presence of ATP. Subsequently, the DNA-protein construct is flushed into the single-molecule flow cell, tethered, and moved into the buffer channel for confocal scanning. **(b)** (left panel) Spatial distribution of fluorescently labeled H2A on DNA containing a chromatinized ARS1 origin as described in **Figure 1b**, deduced from the blue diffraction-limited spots ($N_{foci}$) collected from a number of DNA molecules indicated by $N_{DNA}$. Dashed lines indicate the location of the NPSs, and the solid curve indicates the kernel density estimation of the data. (center panel) Stoichiometry distribution of fluorescently labeled H2A histones in the bin containing the chromatinized origin. Error bars are one-sigma Wilson confidence intervals. White numbers inside the bars designate the fitted values based on the model described in the Methods and in **Supplementary Figure 1.3**. (right panel) Landscape of the mean squared error (MSE) between fit and experiment, calculated for different values of $p_{h2a}$ and $p_{occupancy}$ at a fixed value of $p_{label}$ = 0.81. The red dot indicates the optimal solution with the lowest error. Data derived from two chromatinized samples. **(c)** (left panel) Spatial distribution of fluorescently labeled H2A on DNA containing a chromatinized mutated origin as described in panel (b). (center panel) Stoichiometry distribution of fluorescently labeled H2A histones in the bin containing the chromatinized mutated origin. (right panel) Landscape of the mean squared error (MSE) between fit and experiment, calculated as in panel (b). Data derived from two chromatinized samples.
